# Mountain pine beetle spread in forests with varying host resistance

**DOI:** 10.1101/2024.01.17.575462

**Authors:** Micah Brush, Mark A. Lewis

## Abstract

In the last few decades, mountain pine beetle (MPB) have spread into novel regions in Canada. An important aspect seldom captured in models of MPB spread is host resistance. Lodgepole pine, the predominant host of MPB, varies in resistance across the landscape. There is evidence for a genetic component of resistance, as well as evidence that hosts in areas where MPB has not been present historically are at risk of increased susceptibility. In addition to the spatially varying resistance of the primary host species, the eastward spread of MPB has brought them into jack pine forests. Host resistance in jack pine remains uncertain, but experiments indicate jack pine could be a suitable host. We develop a model of pine beetle spread that links pine beetle population dynamics and forest structure and resistance. We find that beetle outbreaks in the model are characterized by large transient outbreaks that move through the forest. We show how the speed of these outbreaks changes with host resistance and find that biologically plausible values for host resistance are able to stop the wave from advancing. We also find that near the threshold of resistance where the wave is able to advance, small changes in host resistance dramatically decrease the severity of the outbreak. These results indicate that planting trees selected for higher MPB resistance on the landscape may be able to slow or even stop the local spread of MPB. In terms of further eastward spread, our results indicate future outbreaks may move more quickly and be more severe if novel lodgepole pine hosts are indeed more susceptible to beetle attacks, although more research is needed into the susceptibility of jack pine.

## 1 Introduction

The mountain pine beetle (MPB, *Dendroctonus ponderosae* Hopkins) is a bark boring insect native to Western North America (Safranyik and Wilson 2006). MPB have a one year life cycle, emerging in the summer and dispersing to infest hosts (predominately lodgepole pine (*Pinus contorta*). The female beetles bore egg galleries into the phloem of the tree, and the larvae overwinter in the tree before emerging in the following summer. Infested trees generally die the following year. MPB largely maintain small endemic populations, but occasionally erupt in large, landscape scale outbreaks. In western Canada, the most recent outbreak has had severe economic impacts (Corbett et al. 2016; Dhar et al. 2016) and has led to climate change facilitated spread into novel habitats, in particular over the Rocky Mountains and into Alberta (Carroll et al. 2004; Kurz et al. 2008; Sambaraju et al. 2012; Cooke and Carroll 2017; Bleiker 2019). While cold winters have slowed the spread of the beetle for now, there is still uncertainty about the risk of further eastward spread into the boreal forest of Canada (Safranyik et al. 2010; Cullingham et al. 2011; Bleiker 2019).

There are many models, both statistical and mechanistic, that aim to predict the spread of pine beetle (eg. Heavilin and Powell 2008; Powell and Bentz 2009; Strohm et al. 2013; Powell and Bentz 2014; Duncan et al. 2015; Goodsman et al. 2016; Cooke and Carroll 2017; Duncan and Powell 2017; Kunegel-Lion and Lewis 2020; Ramazi et al. 2021; Srivastava and Carroll 2023). One variable that is seldom accounted for explicitly in these models is host resistance (but see eg. Lewis et al. 2010), particularly variation over space. Lodgepole pine, the primary host of MPB, have developed defenses against MPB attack, including the production of a toxic resin to push MPB out of attack holes (Safranyik and Wilson 2006). Beetles can overcome these defenses by mass attacking in large numbers (Raffa and Berryman 1983). The ability of the tree to defend itself against beetle mass attack can vary substantially depending on factors including weather, host vigor, host size, and more (Waring and Pitman 1985; Safranyik and Wilson 2006; Lewis et al. 2010). There is also evidence for a genetic basis for resistance in lodgepole pine (Yanchuk et al. 2008; Erbilgin et al. 2017b; Six et al. 2018), and variation across the range, in particular in areas where MPB was not present historically (Cudmore et al. 2010). Because of the genetic basis for resistance, breeding programs may selectively breed lodgepole pine for MPB resistance, which could lead to more resistant planted cut blocks in the future. In addition to variation in host resistance within lodgepole pine, MPB have expanded their range into jack pine (*Pinus banksiana*) forests in eastern Alberta (Safranyik et al. 2010; Cullingham et al. 2011; Burns et al. 2019). Jack pine is a novel host for MPB, and its susceptibility to MPB attack is uncertain (Cerezke 1995; Safranyik et al. 2010; Rosenberger et al. 2017b; Erbilgin et al. 2017a; Bleiker et al. 2023). More modelling work is needed to understand how this variation in susceptibility across the range, and how future efforts at planting genetically resistant trees, could affect the spread of MPB.

A recent model that couples biologically grounded mountain pine beetle dynamics with forest growth includes explicit parameters for the threshold number of beetles needed to overcome tree defenses as well as the size of brood of MPB in successfully infested trees (Brush and Lewis 2023). The model finds that the population dynamics of the beetles are determined by the ratio of these parameters. The threshold parameter can be easily interpreted in terms of hosts ability to defend itself against mass attack and gives rise to an Allee effect (Goodsman et al. 2016). The brood size parameter can also depend on host quality, where, for example, larger trees produce more beetles (Safranyik and Wilson 2006). Overall then, we interpret the ratio of these terms as the resistance of the host against MPB attack.

We note that here, we refer to host resistance as the ability of an individual pine host to defend against MPB attack. We use the term host susceptibility to indicate both how likely the host is to succumb to infestation more broadly, which includes beetle host preference and host resistance. We make the distinction between host resistance and forest resilience, which we here use to refer to the ability of the forest to recover from MPB outbreaks. Host resistance is therefore one component of forest resilience, and the primary focus of this study.

The goal of the model we develop here is to address questions about MPB spread at the landscape scale with varying host resistance. For example, given evidence for the genetic basis for host resistance in lodgepole pine (Yanchuk et al. 2008; Cudmore et al. 2010; Erbilgin et al. 2017b; Six et al. 2018), is it possible to plant high resistance stands from breeding programs in order to slow or even stop the local spread of a MPB outbreak and thereby give managers more time to respond? Or, given the uncertainty about the susceptibility of novel host species such as jack pine (Cerezke 1995; Safranyik et al. 2010; Rosenberger et al. 2017b; Erbilgin et al. 2017a; Bleiker et al. 2023), what are the risks of outbreak and how might outbreaks be different as MPB continues to move into novel habitat? These types of questions are challenging to answer with existing data or experiments given that they are about events that have not yet occurred, and that MPB outbreaks are unpredictable and depend on environmental and climatic factors. The model we develop here can address some of these uncertainties by simulating MPB outbreaks across a wide range of biological plausible parameter ranges and investigating the qualitative differences in outbreaks across host resistance.

In this study, we extend the model of Brush and Lewis (2023) to include space explicitly in order to study the spread of MPB in hosts of varying resistance. We carefully relate the parameters in this model to MPB data in order to determine a biologically plausible range of host resistance, with realistic levels of spatial aggregation and beetle dispersal. We then investigate how the speed and severity of spread change with host resistance, and how these results may impact further spread at the local landscape scale.

## 2 Methods

### 2.1 Model summary

The model we use to investigate pine beetle spread over time in varying host resistance extends a recently developed model (Brush and Lewis 2023) to include space explicitly. The model is mechanistic and biologically grounded. It includes important aspects of mountain pine beetle biology, modelled after Goodsman et al. (2016), and couples their population dynamics with forest growth, modelled after Duncan et al. (2015). The model dynamics include a strong Allee effect, beetle aggregation over susceptible trees, and a threshold number of beetles for mass attack to overcome tree defenses. The forest structure assumes that new growth is light limited, that beetles do not attack juvenile trees, and that trees have no mortality from causes other than beetle attacks.

To summarize the model, in each year *t* we divide the juvenile trees into *N* age classes, where the number of adult shade equivalents of juvenile trees in each age class *i* is *j*_*i*,*t*_. The juveniles grow each year with survival fraction *s* until they become susceptible trees *S*_*t*_ after *N* years. These trees are assumed to have negligible natural mortality and remain susceptible trees until they are killed by beetles. We define trees to be susceptible when they produce more beetles than it takes to overcome their defenses. The number of female beetles is given by *B*_*t*_, with *c* as a parameter giving the number of female beetles emerging from each infested tree. The fraction of susceptible trees avoiding beetle infestation each year is then given by the function *F* (*m*_*t*_; *k, φ*), where *m*_*t*_ = *B*_*t*_*/S*_*t*_ is the mean number of beetles per susceptible tree, *k* is a parameter indicating the strength of beetle aggregation, and *φ* is a parameter indicating the threshold number of beetles needed to overcome tree defenses. Finally, the number of infested trees is given by *I*_*t*_. We assume these trees lose their needles over two years, where the number of red top trees is *I*_*t-*1_ and the number of gray snags is *I*_*t-*2_. The total number of trees in the fall during census is given by

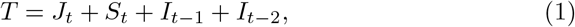

where green infested trees are counted in the *S*_*t*_ class until the following year.

The full evolution of the model from year to year is given by

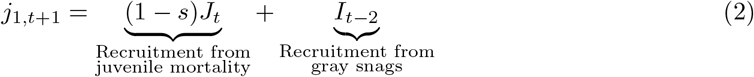

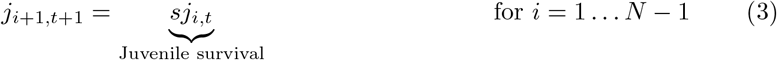

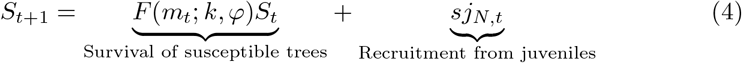

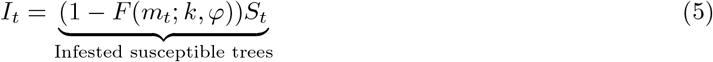

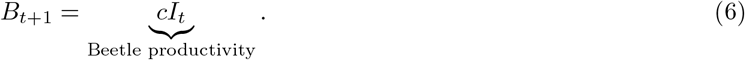

The beetles are assumed to be distributed among the susceptible trees according to a negative binomial distribution with aggregation parameter *k*, and the threshold number of beetles needed to overcome host defenses is *φ*. The function *F* (*m*_*t*_; *k, φ*) describes the fraction of susceptible trees that avoid infestation in year *t* and can be written explicitly as

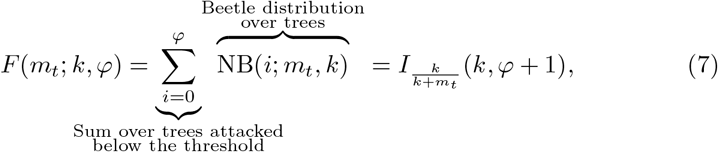

where 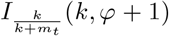 is the cumulative distribution of the negative binomial distribution, which is the regularized incomplete Beta function. The model variables and parameters are summarized in Table 1. More details on the model formulation and the analysis of its structure can be found in Brush and Lewis (2023).

**Table 1:**
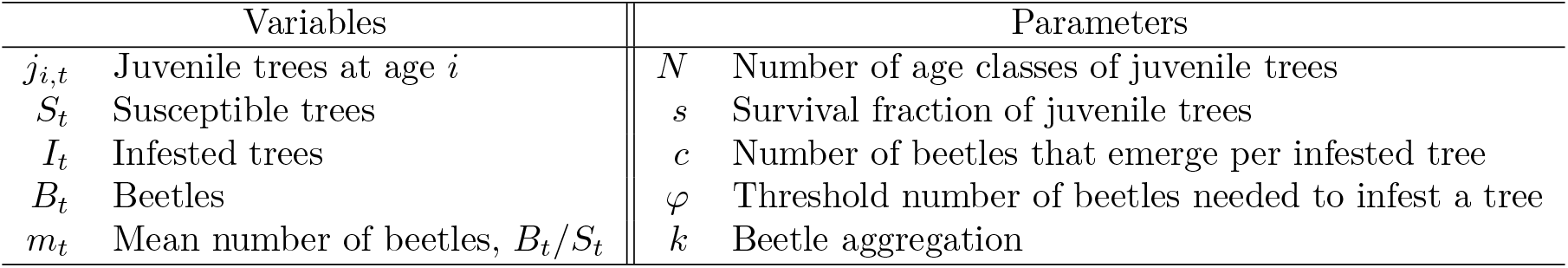
Model variables and parameters.

Mathematically, host resistance in this model is defined by the threshold parameter *φ*. Given that the dynamics of this model depend primarily on the ratio of *φ/c* rather than on either value specifically (Brush and Lewis 2023), we additionally define the ratio *φ/c* as the resistance per tree brood. That this ratio affects the dynamics tells us that the host resistance with respect to the brood productivity parameter is more important for the dynamics than the absolute number of beetles.

### 2.2 Model extension to include space

We now extend the model to include space explicitly, similar to both Goodsman et al. (2016) and Duncan and Powell (2017). We will use integrodifference equations, which are discrete in time but continuous in space (Lutscher 2019). This choice is appropriate given the life cycle and dispersal of the mountain pine beetle (Goodsman et al. 2016; Duncan and Powell 2017; Brush and Lewis 2023). We assume that the beetles disperse each year as they search for new trees to infest with a dispersal kernel *K*, which we discuss in more detail in the next section. We then make each variable dependent on a spatial location *x*. The spatially extended model is then

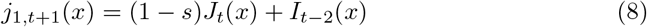

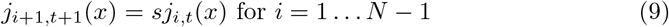

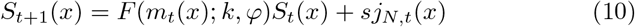

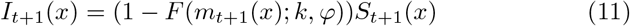

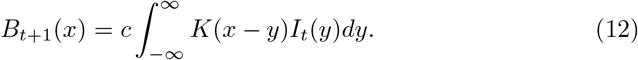

Note that at each spatial location, we still conserve the total number of trees, so *T* = *J*_*t*_(*x*) + *S*_*t*_(*x*) + *I*_*t-*1_(*x*) + *I*_*t-*2_(*x*).

While this extension is theoretically continuous in space, we will consider the case where space is discretized. In practice, this is necessary numerically, but we also interpret the discrete grid biologically. This means the continuous equations are really a continuous approximation of a discrete system. We interpret the discrete grid as individual forest stands with a total number of trees *T*. Within each grid cell the beetles aggregate and infest a fraction 1 *-F* (*m*_*t*_) of the susceptible trees within that stand. We assume that beetles disperse between grid cells, but do not aggregate. Thus, the grid spacing is representative of the scale of aggregation for the beetles.

### 2.3 Beetle dispersal

Mountain pine beetle dispersal is challenging to accurately model, and models quickly become complicated (Berryman et al. 1989; Safranyik et al. 1989; Aukema et al. 2008; Strohm et al. 2013; Powell and Bentz 2014). Beetles largely remain within a few hundred meters (Safranyik et al. 1992; Robertson et al. 2007; Kautz et al. 2011; Carroll et al. 2017), but can also ride convective currents en masse above the tree canopy, where they are taken long distances (tens to hundreds of kilometers) by wind (Furniss and Furniss 1972; Jackson et al. 2008). Both short distance and long distance dispersal can move significant numbers of beetles at the landscape scale (Robertson et al. 2009; Chen and Walton 2011), and there is evidence that beetle dispersal distance depends on stand density (Powell and Bentz 2014). Importantly, short distance dispersal is mediated through complex chemical signalling, where beetles are attracted by host tree semiochemicals, and release pheromones to aggregate for successful mass attack as well as to deaggregate to prevent overcrowding a single tree (Raffa and Berryman 1983; Safranyik et al. 2010; Progar et al. 2014).

Rather than a complex model incorporating wind, chemical signalling, and both short and long distance modes of transportation, we assume that beetle aggregation is accurately captured by the within-cell aggregation model (Eq. 7) and that beyond this scale the beetles follow a dispersal kernel *K*. This model can therefore accurately capture local dispersal, but does not reflect the rare long distance dispersal events.

In past simulations, 2D dispersal kernels are often fit to empirical infestation data (Heavilin and Powell 2008; Goodsman et al. 2016; Goodsman et al. 2017). For mathematical and computational simplicity, as well as to facilitate the interpretation of our model, we will simulate beetle spread in one dimension. This is as if we consider a planar wave in two dimensions, which is an approximation of the case where we have a radially expanding infestation as long as the radius is large enough that the curvature is small locally. To parameterize our simulations, we relate data from beetle dispersal in two dimensions to our one dimensional model using the methods found in Lewis et al. (2016).

We first determine the functional form of the dispersal kernel. In two dimensions, Goodsman et al. (2016), which has the same functional form for the beetle response function as in our model, find that a 2D diffusion and settling kernel outperformed a Gaussian kernel. This kernel can be derived by assuming that beetles disperse as a random walk but drop out onto trees at a constant rate (Broadbent and Kendall 1953; Van Kirk 1995), which is biologically plausible for beetle dispersal at the local scale provided the wind is relatively weak (Safranyik and Wilson 2006). This kernel can be written in two dimensions as

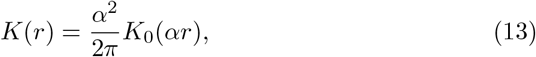

where *K*_0_ is the modified Bessel function of the second kind and zeroth order and *α* is a parameter. That same model of random walk dispersal with constant settling time in one dimension gives the Laplace distribution kernel,

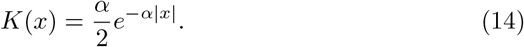

To relate the two dimensional planar case to the one dimensional case, we have to marginalize the 2D dispersal kernel in the direction of wave advance. It turns out that the marginalized probability distribution of Eq. 13 is in fact Eq. 14 (Lewis et al. 2016), and so the *α* parameter in the 1D kernel is the same parameter as that in the 2D kernel that we will use to relate to data.

There is an additional step to connect to data. In 2D, the available data is often the distribution of beetles over radii rather than the full density function. We denote this distribution as 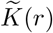, and integrate the full density distribution over all angles, 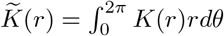 With the kernel in Eq. 13, we find

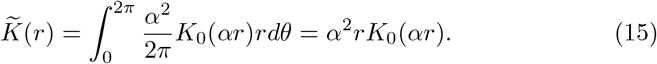

We can now compare this function to data to obtain reasonable values of the parameter *α* that we can then use in our 1D kernel, Eq. 14. Assuming that in our data we find a fraction *p*_1_ of beetles within a radius *r*_1_, we obtain an estimate of α by integrating Eq. 15 out to *r*_1_ and setting that equal to *p*_1_. This integral is equal to

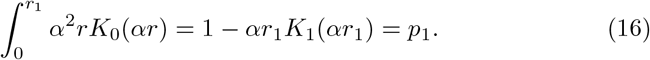

We can solve Eq. 16 numerically to obtain an estimate for α.

We consider three datasets to estimate *α* for our simulations. In a mark recapture experiment, Safranyik et al. (1992) recover 93% and 86% of beetles within 30 meters in two consecutive years (1982 and 1983). Note that these values are the percent of total captured beetles, and so there is likely a bias here where beetles that dispersed beyond the furthest trap at 250 m were not recaptured and are not included in this percentage. Goodsman et al. (2016) use the 93% figure to parameterize their dispersal kernel as they argue that in 1983 MPB had already depleted the tree stock and wanted to parameterize their model assuming a fully stocked stand. Setting *r*_1_ = 30 m and *p*_1_ = 0.93, we find α = 0.121 m^−1^. However, we note that this only takes into account very local dispersal and thus is likely to severely underestimate beetle dispersal. We can also estimate α using results from Robertson et al. (2007). In this study, field crews searched for green attacked trees within 100 m of previous infestations. They find that the the median distance between newly infested stands is 50 meters, assuming newly infested trees came from the nearest previously infested stand. Setting *r*_1_ = 50 m and *p*_1_ = 0.5 we find α = 0.025 m^−1^. Finally, Carroll et al. (2017) use detailed survey data from Alberta to look at infestation spread at a larger scale. They group all infested trees within 750m of each other together into a single parent polygon, which is buffered by 750 m, and then assign all infestations the following year to the closest parent polygon. They find that about 75% of all new infestations occur within 2 km of the nearest parent polygon when averaged across all years from 2007 to 2013. Setting *r*_1_ = 2750 (including the buffer around the parent polygon) and *p*_1_ = 0.75, we find *α* = 0.0008 m^−1^.

Across these three datasets, we find orders of magnitude of variation in α. While most beetles appear to travel only very short distances during an epidemic, the beetles that travel farther are able to infest trees. Given that the goal is to model pine beetle spread at the local landscape scale, we set α = 0.001 m^−1^ in our simulations to accurately capture the spread observed in the highly detailed Alberta survey data.

Importantly, we note that the exact choice of α does not change our qualitative results as it corresponds effectively to a change of units. This is because α has units of m^−1^ is always multiplied by a distance. As long as the scale of dispersal is significantly larger than the grid size of our simulation, our results for the speed of spread are scaled by 1*/*α. See Supporting Information A for more information and plots of the size and speed across the range of α values found here.

### 2.4 Beetle aggregation

In our model, beetles aggregate within a discrete grid cell when attacking host trees according to the survival function *F*. Given the construction of the survival function *F*, this means that the beetles distribute themselves amongst the trees in each cell according to a negative binomial distribution with parameter *k*. We will therefore choose grid spacing that corresponds to the scale of beetle aggregation. Thus while the parameter *α* determines the spatial spread of beetles, the parameter *k* measures how beetles aggregate within discrete grid cells.

We will assume that aggregation occurs at very small scales, within 10 to 30 meters. There are multiple lines of evidence for this scale of aggregation. First, visual evidence from small scale spatial maps, for example in Mitchell and Preisler (1991). Second, Thistle et al. (2004) find that surrogate pheromones disperse to within about 10% of their initial concentration within 10 meters and to only a few percent of their initial concentration within 30 meters. Third, Strohm et al. (2013) build a complex model for beetle dispersal that directly models aggregation pheromones. They use values for their diffusion and degradation parameters from Biesinger et al. (2000), which were derived from dimensional arguments to force the scale of loss of the pheromone to be about 3 m, which is approximately the distance between trees in an open stand. As derived in Chapter 5.2 in Lewis et al. (2016), the dispersal parameter *α* for a kernel with diffusion *D* and constant settling *a* is equal to 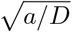, which with the values here is equal to 0.17 1/m. This corresponds to about 92% of aggregation pheromones remaining within 20 m. Finally, this scale of aggregation is consistent with the assumption in the more complicated mechanistic model of Powell and Bentz (2014), which is fit to aerial detection survey data. Here, we will use a grid spacing of Δ*x* = 16 m. We note that typical stands of susceptible lodgepole pine have densities of roughly 1000 stems/ha (Safranyik and Wilson 2006; Boone et al. 2011; Cooke and Carroll 2017), and so Δ*x* =16 m corresponds to approximately 30 stems within a cell, though this could be lower in stands with mixed age structures and lower densities.

The aggregation parameter *k* itself is more challenging to connect directly to data. It is likely that beetle aggregation changes over the course of an outbreak (Safranyik and Wilson 2006; Robertson et al. 2009; Boone et al. 2011; Chen and Walton 2011) or with stand density (Powell and Bentz 2014). Fitting directly to data from Safranyik (1968), Peterman (1974), and Waring and Pitman (1985), we find best fit *k* values ranging from 1.6 to 100, with most values between *k* = 4 and 10. Qualitatively, we find that the negative binomial distribution describes the beetles distribution over trees fairly well, though we note that in some of these cases we fit a zero-inflated negative binomial to capture the many trees with negligible attack densities. We also note that the *k* values that we fit to data describe distributions where beetle populations are large enough to warrant measurement, and thus we would expect these beetles to be less aggregated than in very small populations. We note that part of the reason for the long tail for large *k* is that the distribution changes relatively slowly for large *k* and so the likelihood surface is relatively flat. More information on the data and plots of the fits can be found in Supporting Information B.

Given this variation in *k*, we do not assume a fixed value of *k* and instead assume that beetles at epidemic levels are excellent aggregators at the small scale over which they can detect pheromones. We assume this only for population densities of beetles at the epidemic level as beetle behaviour changes significantly at lower densities, where beetles attack weakened trees in lower numbers and do not rely on aggregation pheromones (Safranyik and Wilson 2006; Boone et al. 2011; Carroll et al. 2017). At epidemic levels, we allow the beetles to aggregate optimally for each value of beetle density. Specifically this means that for any given beetle density the aggregation parameter *k* is chosen so as to maximize the fraction of trees that succumb to infestation. This can also be interpreted as minimizing the fraction of trees that survive infestation. This is a reasonable assumption given the degree of chemical signalling that beetles use to coordinate mass attack at epidemic levels (Raffa and Berryman 1983; Safranyik et al. 2010; Progar et al. 2014). Given that the survival function *F* depends on the mean number of beetles *m*, this also prevents unrealistic behaviour when the number of beetles stays constant but the number of susceptible trees in a stand increases. Without scaling the aggregation, this would mean that the same number of beetles would be less able to infest a more susceptible stand.

Mathematically, we minimize the survival function *F* (*m*; *k, φ*) with respect to *k* for each value of *m*. We set a minimum value of *k, k*_*min*_, where for small values of *m* below epidemic population densities the aggregation is held fixed. We set the epidemic population density threshold according to data from Boone et al. (2011), whose definitions are based on Safranyik and Wilson (2006). They define epidemic population densities as those where more than 20 trees/ha are attacked, and find an average stand density of 1225 tree/ha in their selected stands. Assuming that attacked trees are attacked by exactly the threshold number of beetles needed to overcome defense *φ*, this corresponds to 20*φ* beetles/ha and 20*φ/*1225 beetles/tree in the stand. If we fix *k*_*min*_ at this threshold value of *m*, we find that *k*_*min*_ *≈* 0.007 to 0.008 across the range of *φ* considered. We find that this number increases if beetles attack in numbers above the exact threshold *φ*, and so round up to *k*_*min*_ = 0.01 in our simulations. We note that changing *k*_*min*_ does have an effect on the Allee threshold at low beetle densities, which makes a difference for when beetles are able to infest new stands. The effects of *k*_*min*_ are studied further in Supporting Information C.

For very large values of beetle density, *m> φ*, this optimization would result in *k → ∞* as the beetles would ideally spread completely uniformly across all trees. We assume there is some maximum value of *k, k*_*max*_, so that the beetles will always aggregate to some extent even if their density is very high. Biologically, this reflects that the aggregation pheromones are present before the anti-aggregation pheromones (Progar et al. 2014). We set *k*_*max*_ = 100 in simulations. We note that our results are not that sensitive to the exact value *k*_*max*_ above about 50 when the negative binomial becomes well approximated by the Poisson distribution. The effects of *k*_*max*_ are also studied further in Supporting Information C.

We plot this optimized *F*, as well as the corresponding values of *k* that minimize the function for each value of *m*, in Fig. 1. We additionally plot the Allee threshold in terms of mean number of beetles per susceptible tree in Fig. 2. Numerically, we optimize *F* by minimizing the function with respect to *k* for each integer value of *m* between a small value (chosen to be 11) and the threshold *φ* plus 100. For *m* below this value, we optimize for logarithmically spaced values between 0.01 and 10. For values of *c* greater than *φ*+ 100, we also sample logarithmically up to *m* = *c*. We choose this range as for very small or large *m*, the function *F* is relatively insensitive to *k* and optimization becomes difficult numerically. Below and above this range of *m*, we set *F* = 0 and *F* = 1, respectively. Within this range we interpolate the function between the values of *m* where we optimized. When *k*_*min*_ and *k*_*max*_ are fixed, we set the range of *k* directly when optimizing. When *k*_*min*_ is instead fixed given a threshold value of *m*, we set the minimum *k* during optimization to 10^*—*4^ and then afterwards find the value of *k* at the threshold value of *m* to use as *k*_*min*_.

**Figure 1.**
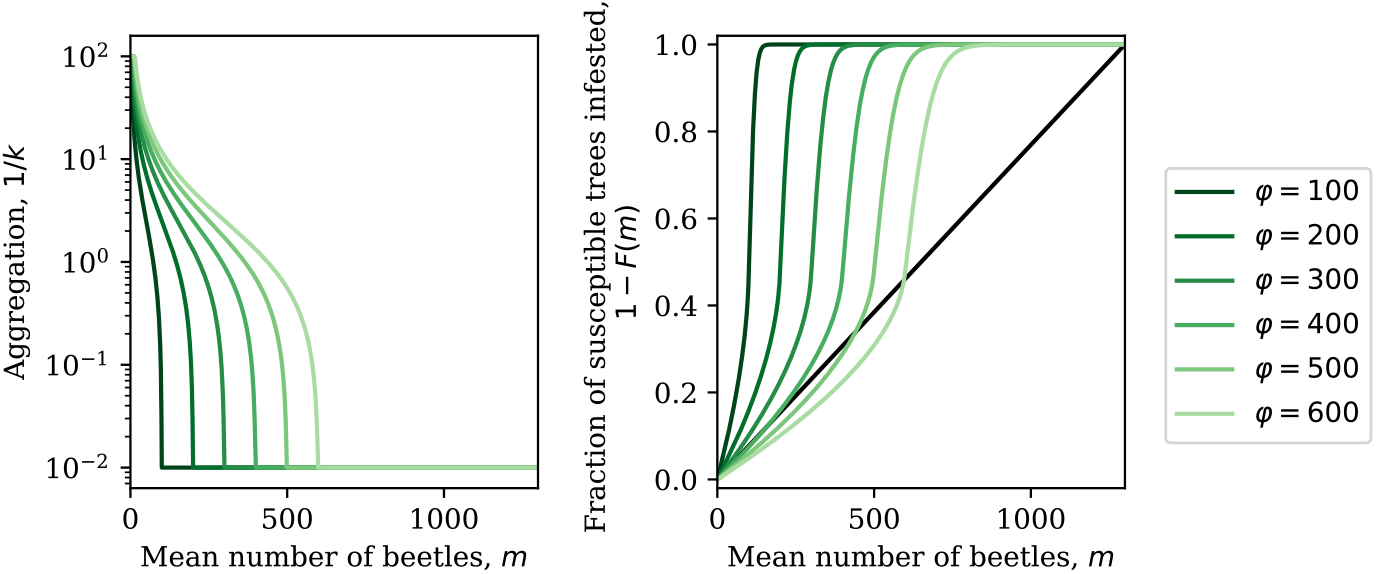
Optimized aggregation across mean beetle density and the corresponding function 1 − *F* (*m*) representing the fraction of susceptible trees infested for different values of the threshold *φ* and *c* = 1300. We set the bounds for *k* to be *k*_min_ = 0.01 and *k*_max_ = 100. These cutoffs are visible in the plot of 1*/k*.

**Figure 2.**
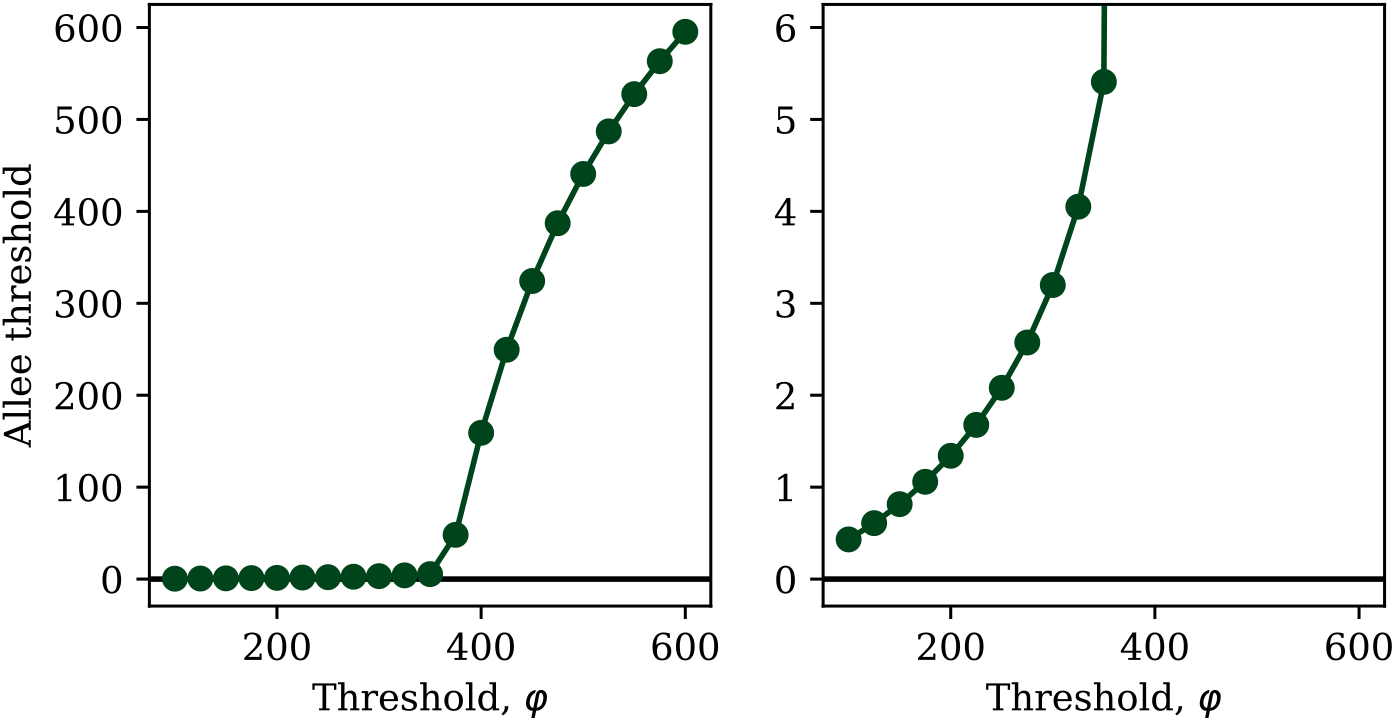
The Allee threshold at different values of the threshold parameter *φ* with *c* = 1300, given *k*_min_ = 0.01 and *k*_max_ = 100. The Allee threshold is given in terms of the mean number of beetles per susceptible tree. The second panel shows the same information but with a different *y*-axis to show smaller values of the Allee threshold more clearly.

### 2.5 Parameters for population dynamics

The parameters for population dynamics in the model are the number of years before trees become susceptible *N*, the juvenile survival *s*, the brood size parameter *c*, and the threshold parameter *φ*. We find plausible values for each of these parameters from available data in the literature for MPB in lodgepole pine. Table 2 summarizes the range we find for each parameter, and the single value used in simulations with fixed parameter values. We here summarize our parameter estimates and provide more details in Supporting Information D.

**Table 2:**
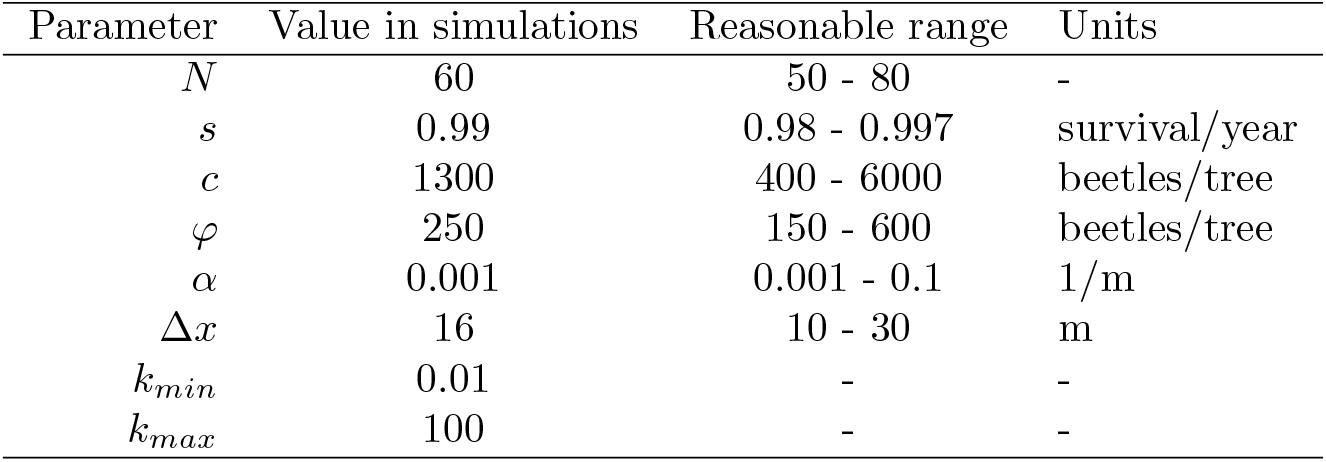
Values of model parameters used in numerical simulations, along with reasonable ranges from data.

| Parameter | Value in simulations | Reasonable range | Units |
| --- | --- | --- | --- |
| $N$ | 60 | 50 - 80 | - |
| $s$ | 0.99 | 0.98 - 0.997 | survival/year |
| $c$ | 1300 | 400 - 6000 | beetles/tree |
| $\varphi$ | 250 | 150 - 600 | beetles/tree |
| $\alpha$ | 0.001 | 0.001 - 0.1 | 1/m |
| $\Delta x$ | 16 | 10 - 30 | m |
| $k_{min}$ | 0.01 | - | - |
| $k_{max}$ | 100 | - | - |

We estimate the number of age classes of juvenile trees *N* using two methods. In both cases, we note that because we assume brood size is the same across all susceptible trees, we define susceptible trees as those that produce more beetles than it takes to overcome their defenses. This means that the number of age classes of juvenile trees will be slightly larger than if we were to estimate only the number of years before trees could be attacked at all. First, in some outbreak data, tree age was recorded and we use that directly. In these data, we find that trees become susceptible around age 55 to 60 (Safranyik 1968; Safranyik and Wilson 2006; Shore and Safranyik 1992). Second, we estimate the diameter at breast height (dbh) cutoff for when trees become susceptible and combine that with estimates of growth. From data, we find the dbh cutoff varies between 18 cm and 35 cm (Klein et al. 1978; Mitchell and Preisler 1991; Safranyik and Wilson 2006; Safranyik et al. 1974; Cole and Amman 1969). We set the cutoff at 25 cm. We then use two approaches for growth estimates. First, we use radial growth rates, which vary between 1.2 mm/year and 2.4 mm/year (Chhin et al. 2008; McLane et al. 2011; Heath and Alfaro 1990). Rounding these to 1.5 mm/year to 2.0 mm/year with a dbh cutoff of 25 cm corresponds to *N* = 63 to 83. Given that growth rates are highly dependent on location, we also use site index to estimate growth, which is defined to be the height the average tree will reach 50 years after reaching breast height and is mapped throughout BC and Alberta. Typical site indices in Alberta are 18 to 21 m (Monserud et al. 2008; Thrower et al. 1994), which translate to 25 cm dbh using a height to dbh model (Huang 1999). We then add the 6 years it takes to reach breast height (Huang et al. 1994; Thrower et al. 1994; Nigh and Love 1999) to obtain an estimate of *N ≈* 56. Combining all estimates, a plausible range is from *N* = 50 to 80 and we set *N* = 60 in simulations.

We estimate the survival fraction of juvenile trees *s* directly from juvenile survival data. We note that in reality juvenile mortality is higher for younger trees, though here we assume it is the same across all age classes. This means that this parameter will be larger than from data for 5 or 10 year survival rates. Estimates range from *s* = 0.94 to *s* = 0.997 (Brown and Navratil 1995; Bedford and Sutton 2000; Rweyongeza et al. 2007) for 10 year survival rates, and so we find a plausible range from *s* = 0.98 to 0.997 and set *s* = 0.99 in simulations.

Many sources of data for both the brood size per tree *c* and host resistance are measured in terms of beetles per m^2^. To relate to these parameters, we have to estimate the surface area of a tree attacked by beetle. Previous estimates range between 4.4 and 8.1 m^2^ per tree (Strohm et al. 2013; Powell and Bentz 2009; Klein et al. 1978). We average these estimates and round to 6 m^2^ per tree, though note in cases with stand size distributions we make use of that information to obtain more accurate estimates.

We estimate the brood size parameter *c* from several different data sources, noting that this parameter is likely to be highly variable across years and host tree. Additionally, because we model only the population dynamics of the female beetles, we will assume that 2/3 of all beetle offspring are female (Amman and Cole 1983; Safranyik and Wilson 2006). We can estimate the brood size by multiplying the attack density per m^2^, the attacked surface area, the number of offspring per attack, and the sex ratio of the offspring. Estimates using this method range from 400 to 3000 female beetles per tree (Klein et al. 1978; Raffa and Berryman 1983; Goodsman et al. 2016). We can also use direct estimates of emergence data per tree with known dbh, which range from a few hundred to 15 000 beetles, and then average over stand size distributions to obtain an estimate of around 1250 female beetles per tree (Safranyik 1968; Safranyik and Wilson 2006; Cole and Amman 1969; Powell and Bentz 2009). We find a plausible range from from *c* = 400 to 6000 female beetles per tree and set *c* = 1300 in simulations.

We estimate the threshold *φ* from experiments manipulating aggregation at 40 beetles per m^2^ or *φ* = 240 beetles per tree (Raffa and Berryman 1983). We estimate a plausible range of host resistance to be *φ* = 150 to 720 beetles per tree using data from Waring and Pitman (1985) and the fit to that data in Lewis et al. (2010). Note however that the upper range here is only obtained by a few trees, and our parameter is at the stand level. We also obtain estimates between *φ* = 250 and 520 beetles per tree from our fits to data from Peterman (1974) (Supporting Information B). We find a plausible range from *φ* = 150 to 600 and set *φ* = 250 in simulations.

Finally, we note that the qualitative results of this model depend primarily on the ratio of *φ/c*, rather than on either value specifically (Brush and Lewis 2023). The range of *c* we find here varies by a factor of 15, and the range of the threshold *φ* varies by about a factor of 4; however both of these are likely to vary with similar factors (tree size, primarily (Safranyik and Wilson 2006)) and are not independent. Assuming they vary together linearly, the ratio *φ/c* varies between 600/6000 = 0.1 and 150/400 = 0.375. We can only estimate this ratio directly using data from Raffa and Berryman (1983). They find a threshold of 40 beetles per m^2^ and productivity between 200 and 500 beetles per m^2^, corresponding to *φ/c* roughly between 0.08 and 0.2. If we were to vary *φ* across its plausible range from 150 to 600, this gives a similar range for *φ/c* of 0.12 to 0.46. In most plots we will round the plausible range to be from 100 to 600 beetles per tree, which corresponds to *φ/c* between 0.08 and 0.46. This also allows for the possibility that host defense thresholds are lower in jack pine or in novel stands.

### 2.6 Simulations

We simulate the mathematical model defined by Eqs. 8–12 in space with a discretized grid to investigate MPB spread. We set the initial conditions such that half of the forest is already infested by beetles and then simulate the model forward in time to observe the outbreak dynamics. We define metrics to characterize the speed and size of the outbreak and investigate how these change with host resistance.

On a technical level, we implement 1D spatial simulations on a discrete grid with spacing Δ*x* in Python (code available at https://github.com/micbru/MPBSpatialModel/). We perform the convolution in Eq. 12 by taking the product of the infested tree density and the dispersal kernel in Fourier space by taking the discrete Fourier transform (DFT) using Fast Fourier Transforms (FFT), specifically using `scipy.fft` (Lutscher 2019, see for eg.).

We interpret the discrete grid as individual forest stands, as discussed above, where beetles aggregate and infest a fraction 1 − *F* (*m*_*t*_) of the susceptible trees within that stand. Within each stand, the total number of trees is fixed to 1 (*T* = 1 in Eq. 1) so that we interpret susceptible or infested tree density as a fraction of the total trees in the stand. We can then convert from this to the number of stems by assuming a stand density. We note that realistic stand densities for susceptible stands are roughly 1000 stems/ha (Boone et al. 2011; Cooke and Carroll 2017; Safranyik and Wilson 2006), and given Δ*x* = 16 m, this corresponds to about 30 stems/cell. We use a grid much larger than the range shown in plots or used to calculate wave speeds to avoid edge effects.

We initialize our simulations by first optimizing the beetle aggregation for our set of *φ* and *c* as described above, and then setting all variables on the left half of the spatial grid (*x <* 0) to their stable equilibrium values obtained by solving 1 − *F* (*m*) = *m*/*c*. This makes the forest structure geometric. Given that the total number of trees depends on the number of infested trees in the two previous years (Eq. 1), we assume for *x <* 0 that the number of infested trees in the two previous years were also at equilibrium. For *x ≥* 0, we set all trees to be susceptible, *S*_0_(*x ≥* 0) = 1. We assume that no trees were previously infested. This makes it easier to interpret the peak infestations as a fraction of the susceptible trees.

To determine the speed of the wave, we find the furthest point along the wave where the beetle population is greater than half of the equilibrium density at each time, 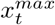, and then take the difference to obtain the number of meters the population moves each year, 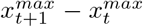. We then average this distance over the second half of the simulation. Note that we also check to ensure the wave does not collide with the backwards travelling wave from edge effects, and if it does we only compute 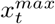 up until the waves collide.

To determine the peak size of the wave, we take the maximum value of the wave at any point in space and average over a set time period. We start the time period at the first time the fraction of infested trees grows above its equilibrium value at *x* = 0 plus half of the period, and average over an entire period.

## 3 Results

We are interested in the characteristics of spread when beetles move between areas of different host resistance *φ*, or equivalently areas of different host resistance per tree brood *φ*/*c*. This could represent spread into a region of more genetically resistant lodgepole pine (Erbilgin et al. 2017b; Six et al. 2018), or into jack pine where host susceptibility is uncertain given host thresholds are likely to be lower but the beetles reproductive potential may also be lower (Safranyik and Linton 1982; Cerezke 1995; Safranyik et al. 2010; Rosenberger et al. 2017a; Rosenberger et al. 2017b; Bleiker et al. 2023).

In numerical simulations across a wide range of values of the threshold *φ* (and correspondingly *φ*/*c*), the infestation advances in space as a travelling transient pulse with beetle population densities much higher than at equilibrium. These travelling pulses are separated in time with period *N* + 3. This is as the transient waves in the non-spatial case in Brush and Lewis (2023), but here the waves are driven explicitly by beetles spreading over space into novel stands. Figure 3 shows a 3D representation of the beetle density over 200 years with the parameter values in Table 2, and Supporting Video A1 shows the same simulation as an animation.

**Figure 3.**
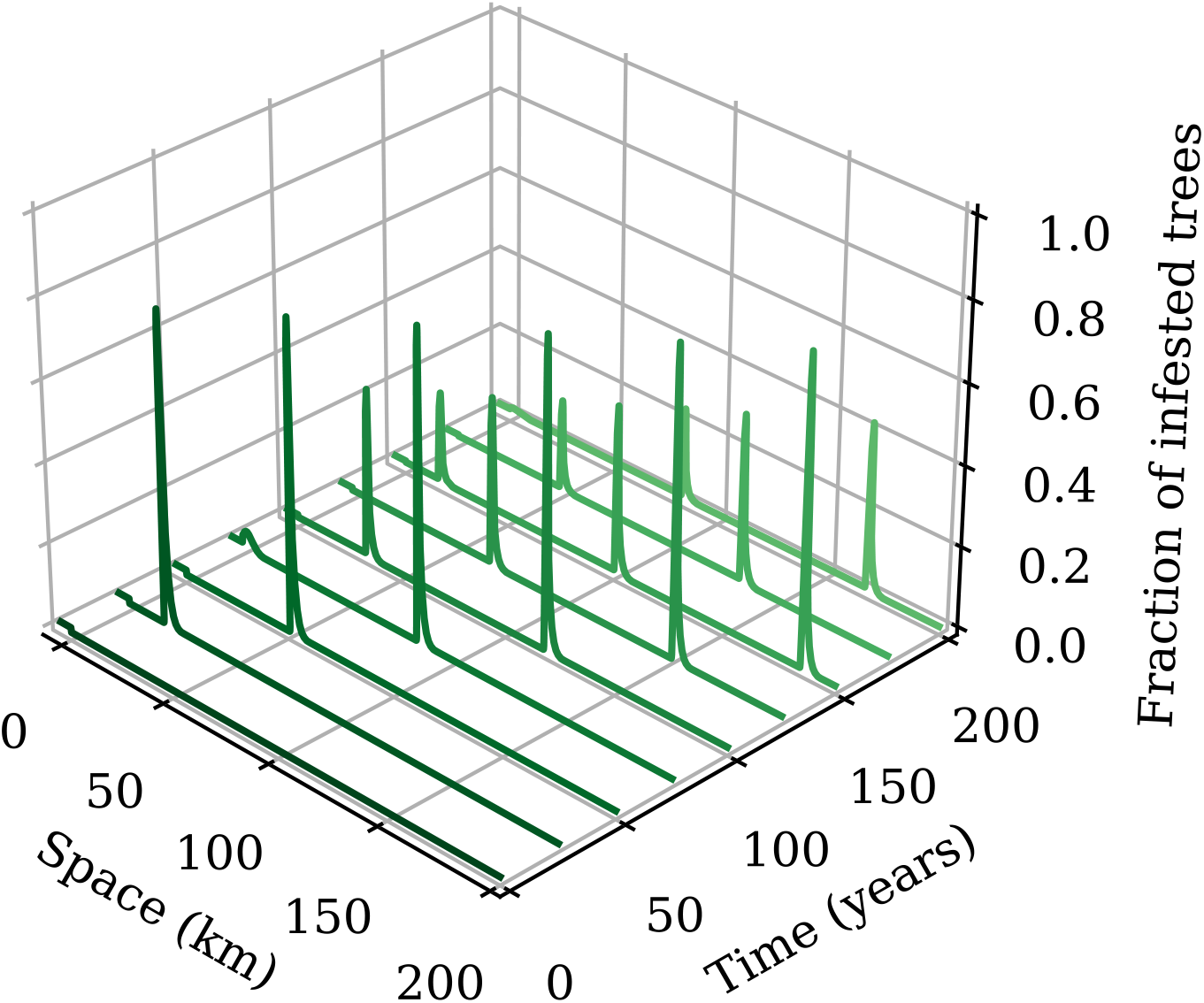
The fraction of infested trees over time and space. We initialize the forest structure and beetle population to their equilibrium values for *x<* 0. For *x ≥* 0, all trees are set to susceptible with no initial beetles. The parameter values are as in Table 2 (*c* = 1300) and we run the simulation for 200 years.

Figure 4 shows the speed of spread for different values of *φ* with the other parameters fixed to their values in Table 2. Again, while we vary only *φ*, since the ratio *φ*/*c* determines the dynamics this is equivalent to holding *φ* fixed and varying *c*, or varying both parameters simultaneously with a specific ratio. The simulations were run for 200 years with increments of 25 for the threshold *φ*. The speed increases nearly linearly with decreasing *φ <* 350 (*φ*/*c <* 0.27), with a sharp change between 325 *< φ <* 375 (0.25 *< φ*/*c <* 0.29) as soon as the wave is able to advance. For *φ >* 350 (*φ*/*c >* 0.27, the wave is unable to advance. Note that these speeds scale with 1/*α*, unless the scale of dispersal is similar to the scale of aggregation (Supporting Information A).

**Figure 4.**
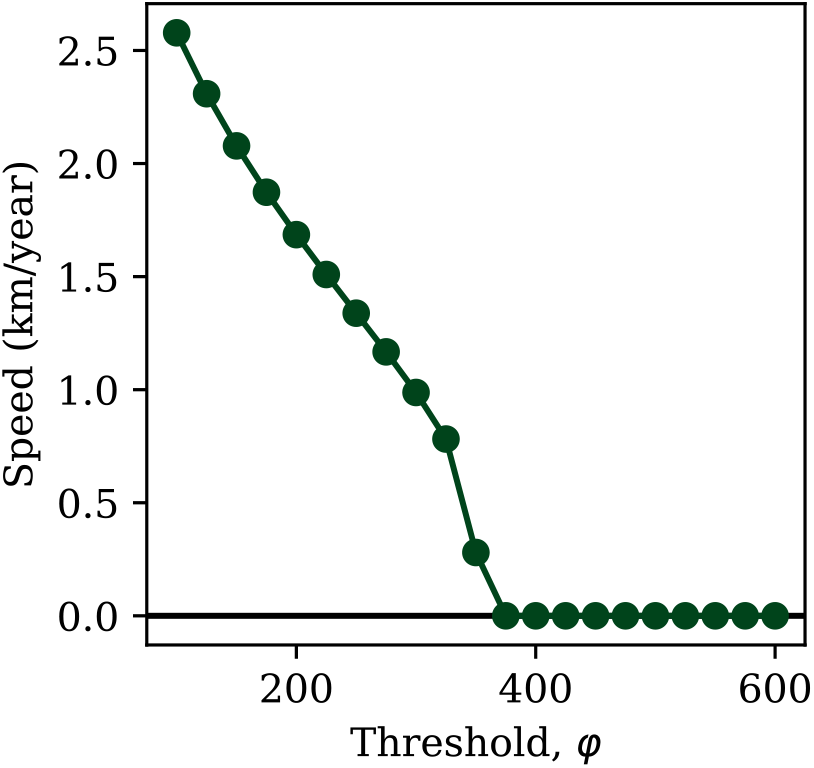
The speed of the transient wave pulse for different values of the host resistance. The other parameter values are as in Table 2 (*c* = 1300) and we run for 200 years. Points indicate values of the threshold where the speed was calculated and the line is the linear interpolation between the points.

Figure 5 shows the averaged peak fraction of infested trees in the transient wave pulse for the same simulation as in Fig. 4. We again note the sharp jump in the size of the infestation when *φ ≈* 350 (*φ*/*c* = 0.27) as the wave is suddenly able to advance. Another potential metric for measuring the size of the outbreak would be total outbreak footprint, or the total number of infested trees over one period in time for one transient outbreak. In the asymptotic case considered here where all trees are susceptible, the total footprint of the outbreak is 1, or all trees are infested over the course of one transient wave. We note that in simulations where the right hand side started at equilibrium forest density rather than with all trees susceptible, we observe very similar dynamics, including speed, but the magnitude of the peak infestation was smaller.

**Figure 5.**
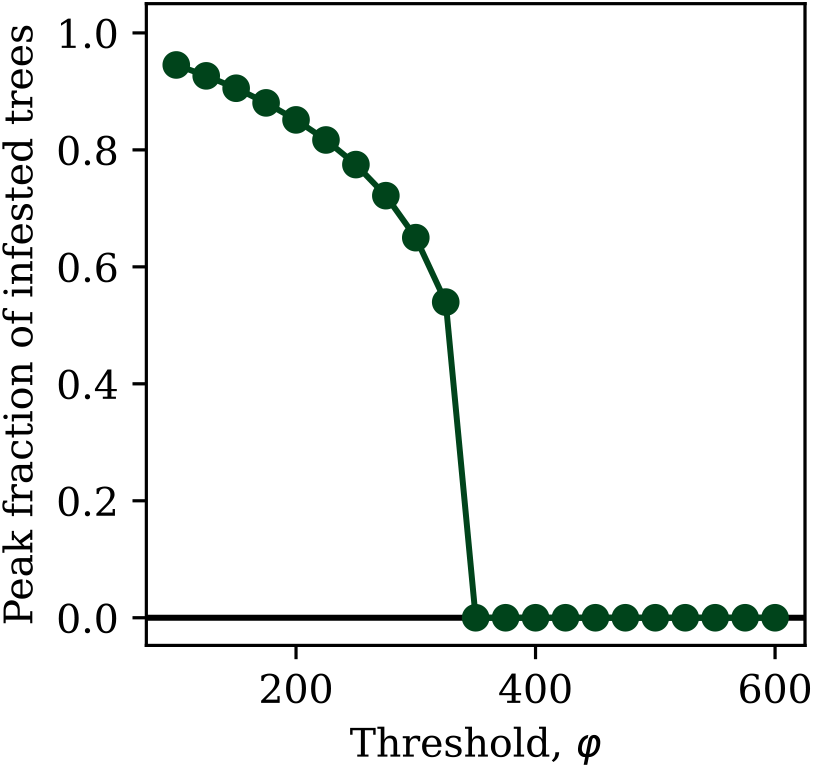
The averaged peak fraction of infested trees in the transient wave pulse for different values of the host resistance. The other parameter values are as in Table 2 (*c* = 1300) and we run the simulation for 200 years. Points indicate values of the threshold where the size was calculated and the line is the linear interpolation between the points.

We now focus on simulations with spatial variation in *φ*. Figure 6 shows how the wave advances from *φ* = 200 to 300 and then 400 (*φ*/*c* = 0.15, 0.23, 0.31), with Fig. 7 showing the dynamics at three fixed points in space in each of the different *φ* region. The results here reflect those in Fig. 4 in that the speed is slower when *φ* = 300 and the wave can no longer advance at *φ* = 400. The same simulation can be viewed as an animation in Supporting Video A2.

**Figure 6.**
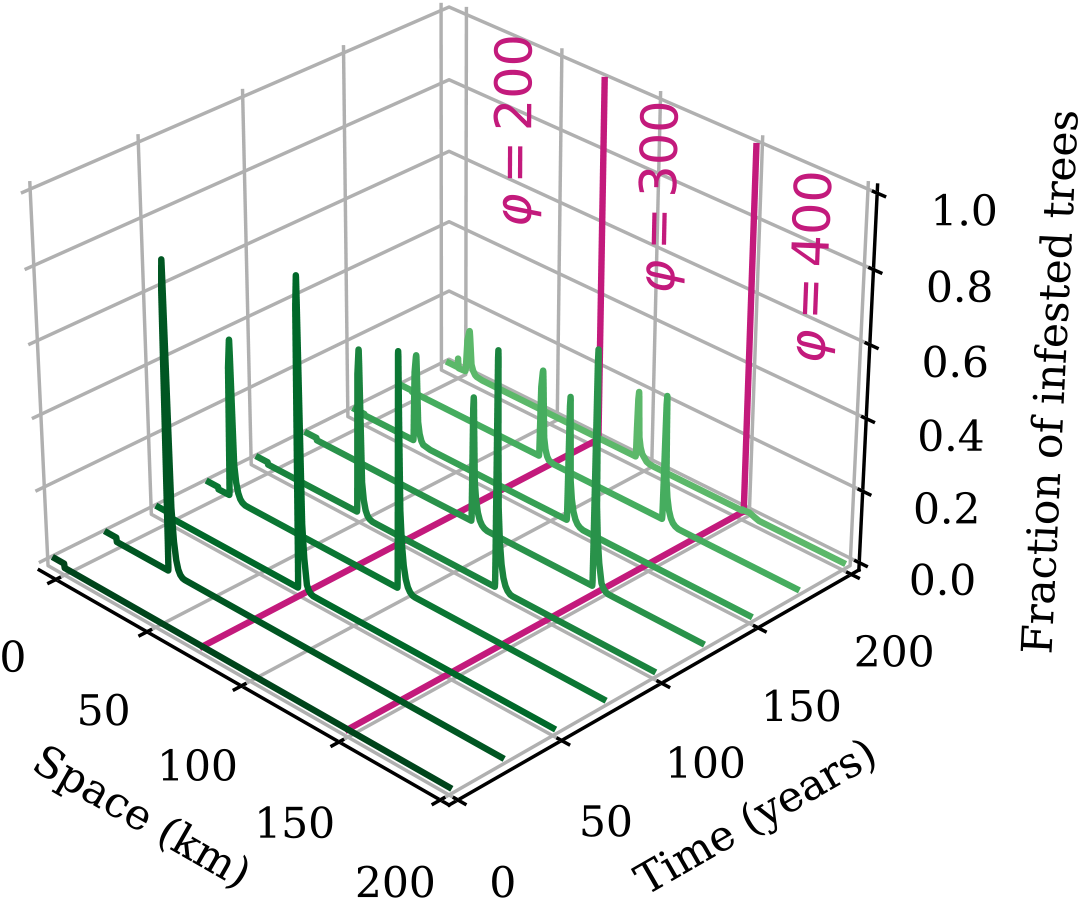
The fraction of infested trees over time and space with varying host threshold. We initialize the forest structure and beetle population to their equilibrium values for *x <* 0. For *x ≥* 0, all trees are set to susceptible with no initial beetles. We set the threshold *φ* to 200, 300, and 400 beetles per tree for *x <* 75 km, 75 km ≤ *x <* 150 km, and *x ≥* 150 km respectively. The other parameter values are as in Table 2 (*c* = 1300) and we run the simulation for 200 years.

**Figure 7.**
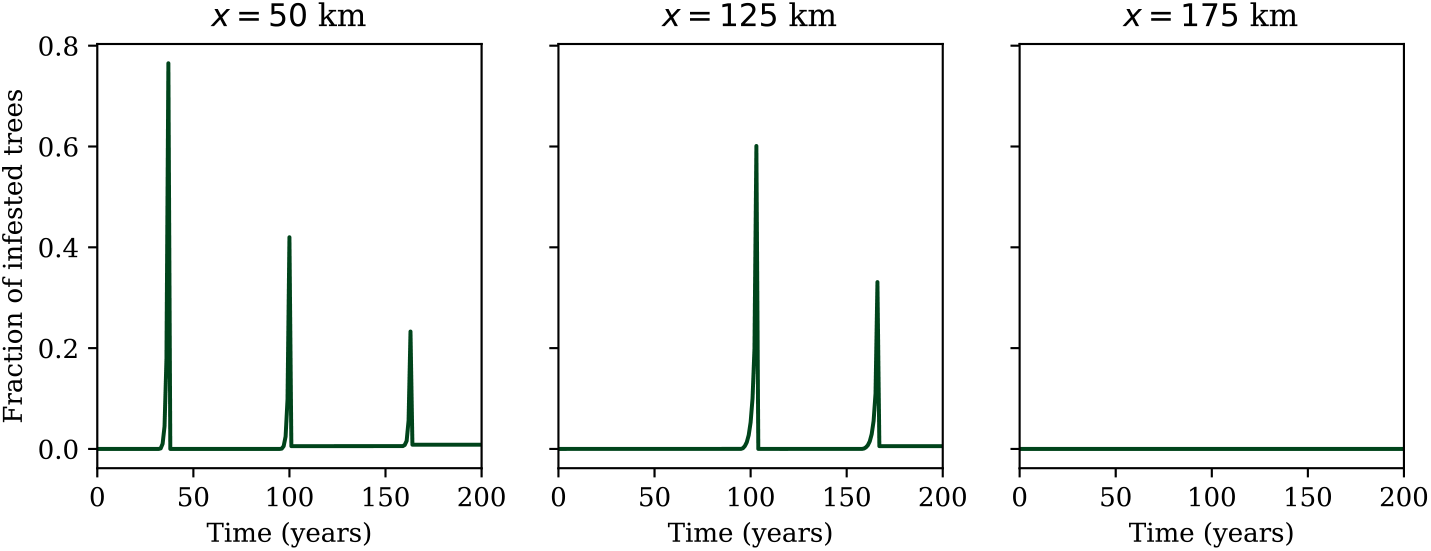
The fraction of infested trees over time with varying host threshold at three different fixed points in space. The simulation is as in Fig. 6. The three spatial locations are chosen to be within each of the different regions of the threshold *φ*.

We also consider the case of a buffer zone where beetles pass through a section of higher resistance hosts. This could represent spread across a stand planted with trees from a breeding program that were selected for host resistance. The buffer section with higher resistance begins at *x* = 0 and extends for widths between 1 and 4 km in increments of 0.5 km. We set the threshold to overcome tree defenses for the buffer zone to be *φ*_*b*_ and vary it from 400 to 600 in increments of 25, with the lower bound chosen to be above the value where the wave would not spread from Fig. 4. In Fig. 8, we show the time it takes for the beetles to pass through the buffer for each combination of width and value of *φ*_*b*_, where the dark hashed region in the upper right indicates the beetles were not able to pass through the buffer zone in the 200 years of the simulation. Supporting Videos A3 and A4 show animations of beetle density for scenarios where the beetle can and cannot spread beyond the buffer.

**Figure 8.**
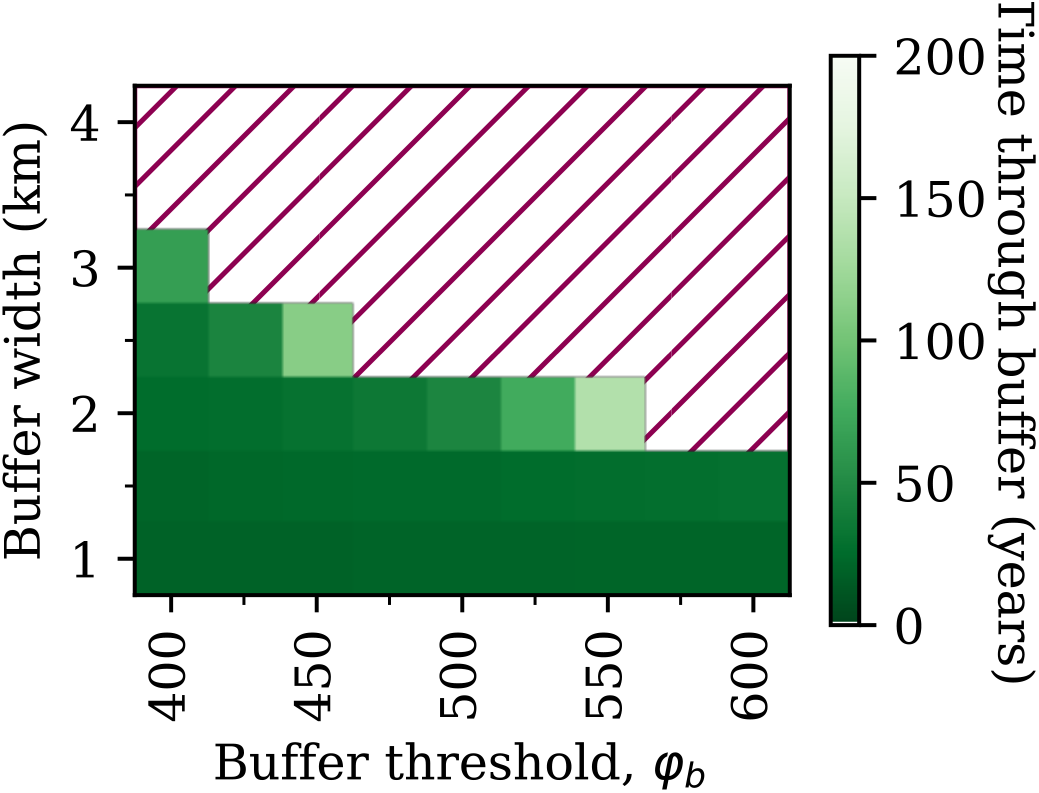
Beetle population spread through a buffer zone with higher host resistance for a variety of buffer widths and values of buffer host resistance. For each combination of parameters, we run the simulation for 200 years. If the beetles are able to pass through the buffer, we color the corresponding cell green with lighter colors indicating that the beetles took a longer time to pass through the buffer. Parameter regions where beetles are not able to pass through the buffer are indicated by dark red hashes. We initialize the forest structure and beetle population to their equilibrium values for *x<* 0. For *x ≥* 0, all trees are set to susceptible with no initial beetles. The buffer itself starts at *x* = 0 with threshold *φ*_b_. All other parameters are as in Table 2 (*c* = 1300).

## 4 Discussion

Our model simulations show that spread of MPB at the landscape scale can be stopped at relatively intermediate values of host resistance (Fig. 4). Below that value when the spread does proceed, the speed of the outbreak depends approximately linearly on host resistance (Fig. 4), but the peak size of the outbreak is comparatively less affected by the host resistance (Fig. 5). Given that the value of host resistance where spread is stopped is within a plausible range of natural resistance, it is possible that planted trees selected for resistance could slow or stop local spread, even if they are not planted everywhere on the landscape (Fig. 8). More research is needed to determine the relevant parameter values in jack pine, but recent research points towards lower brood sizes in jack pine (Bleiker et al. 2023), which could mean spread further east in Alberta will progress more slowly than in the past few decades.

Broadly, the outbreaks in our simulations of spatio-temporal dynamics of MPB dynamics are characterized by large transients that move through the susceptible trees, as in the non-spatial model (Brush and Lewis 2023). These transient waves of beetles are many times larger than the equilibrium beetle density and decay over time. When we view the outbreak from a fixed location in space, as in Fig. 7, the dynamics appear very similar to the non-spatial dynamics in Brush and Lewis (2023).

We find that the beetle population is no longer able to spread at the intermediate value of *φ* = 350 with *c* = 1300. This value of host resistance corresponds to approximately 60 beetle attacks per m^2^ to overcome tree defenses, which is near the intermediate value of host resistance found in both Lewis et al. (2010) and Peterman (1974). This means that a relatively modest increase in host resistance may be able to slow or stop local spread for outbreaks of intermediate size, as this value of host resistance assumes *c* = 1300 beetles per tree. When we look instead at the value of host resistance per tree brood *φ*/*c ≈* 0.27, it is indeed greater than the maximum value of *φ*/*c* obtained from the data in Raffa and Berryman (1983), which is the only dataset for which we can calculate host resistance per brood tree. That this value is larger than the threshold value makes sense as the forest stands under study were undergoing an outbreak. It is unlikely that the host resistance per tree brood will be observed to be above about 0.25 as in that case the speed and size of the wave are very small or zero (Figs. 4 and 5), and such a small outbreak may be undetected or not studied in the level of detail necessary to obtain estimates for this ratio.

We note that the critical value of *φ*/*c ≈* 0.27 where the wave is no longer able to spread is much lower here than the appearance of the upper stable equilibrium at *φ*/*c ≈* 0.6 found in the non-spatial model (Brush and Lewis 2023). This means that *φ*/*c* must be substantially below the criteria for the existence of the upper equilibrium in order for the wave to spread. In the nonspatial model, this equilibrium corresponds to when *φ*/*c* is small enough that infested trees are able to replace themselves. In the spatial model, this value is smaller as beetles disperse out of infested cell and so either the brood size must be larger or the host resistance must be smaller in order for the beetles to replace themselves. A mathematical approximation is that the outbreak is able to spread when the number of beetles in equilibrium divided by two is equal to the Allee threshold. The factor of one half comes from the fact that in our setup, half of the initial space is infested by beetles, and so the dispersing beetles contribute roughly half of the equilibrium number at *x* = 0. Numerically with parameter values from Table 2, we find the equilibrium number of beetles is about 15 beetles/cell with *T* = 1 (which corresponds to about 450 beetles/cell with stand densities of around 1000 stems/ha). Assuming *k ≈ k*_*min*_ near the Allee threshold, we find that the approximate value of *φ* for which the wave advances is *φ* = 366 by solving 1 − *F* (*m*; *k*_*min*_, *φ* + 1) = *m*/*c* with *m* = 15/2 and *c* = 1300, which is roughly consistent with our numerical findings.

From a theoretical perspective, we can obtain an upper bound for the value of *φ*/*c* where the speed is zero and the wave reverses direction by analyzing the model with no forest structure and where infested trees are immediately replaced by susceptible trees (or in other words, infested trees are not killed). The spread in this case is faster as there are always plenty of susceptible trees for the beetles, which is why this represents an upper bound. Mathematically, the model analysis in this case is simplified considerably and we can show that the wave is no longer able to advance when 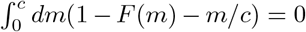 (Wang et al. 2002; Li and Otto 2022). This is true for *φ ≈* 750 with *c* = 1300, or *φ*/*c ≈* 0.58 (see Supporting Information E). This is well beyond the realistic range for *φ*/*c*, and so while transient outbreaks may be slowed or stopped, they are unlikely to be reversed completely as that would require even larger *φ*/*c*.

We can also consider when the wave is able to spread in the context of forest thinning, which has been found to be effective against beetle infestation (Waring and Pitman 1985; Safranyik and Wilson 2006). Typical stand densities in our model correspond to approximately 30 stems/cell and 450 beetles/cell. Thinning the forest decreases this absolute number of beetles by decreasing the number of stems/cell, while also likely increasing the resistance of the trees in the cell by increasing their vigor (Safranyik and Wilson 2006; Waring and Pitman 1985). Even without the increase to resistance, thinning to a density of 500 stems/ha (Safranyik and Wilson 2006) would result in only about 200 total beetles per cell, making it difficult for beetles to overcome even moderate host resistances. Note that this analysis demonstrates where considering the total number of trees per cell *T* does affect the results.

We find that the spread speeds are consistent with data for intermediate length mountain pine beetle spread, or spot proliferation (Robertson et al. 2007; Robertson et al. 2009; Chen and Walton 2011). More specifically, our results for the speed of spread are consistent with the analysis of Robertson et al. (2009), who find that the median displacement from ground spread in three regions on the BC-Alberta border ranges from a few hundred meters to just over a kilometer (where spread over 2 km is classified as long distance dispersal). This is unsurprising given we parameterized our dispersal kernel using data from intermediate distance data from Carroll et al. (2017). We note here that setting *α* according to the results from Safranyik et al. (1992) or Robertson et al. (2007) would be more suitable for within stand dispersal, but that this scale is less relevant for determining the speed of spread as beetles do not move beyond the local stand. Since the speed of dispersal scales as 1*/α*, we find that when the dimensionless parameter Δ*x · α* is greater than about 2, the transient wave can fail to advance due to the discreteness of the grid. This is because the dispersal into the adjacent cell is not enough to overcome the Allee effect, and so the wave is pinned (Keitt et al. 2001). With *α* = 0.001 m^*—*1^, we would only expect to see this pinning if beetles aggregated on the scale of roughly 2 km, which is far from realistic values. Note that this means our results are not that sensitive to the exact choice of Δ*x*, and so a grid spacing of 32 or 64 m (which may be better suited to less dense stands) would not significantly impact our results. See Supporting Information F for more information.

Note that the spread speeds we find here are smaller by about an order of magnitude than the 80 km/year in Alberta found by Cooke and Carroll (2017); however that spread includes long distance spread events, which we do not model explicitly here. These long distance dispersal events happen when beetles are transported by the wind above the tree canopy (Furniss and Furniss 1972; Jackson et al. 2008) and are known to be important for infestations in new areas (Robertson et al. 2009; Chen and Walton 2011), even though a relatively small number of beetles (likely a few percent) disperse in this way (Safranyik et al. 1992). One possibility is to extend this model to include stratified diffusion (Shigesada et al. 1995) and to allow for a small number of beetles to travel above the canopy with a separate mode of dispersal. We additionally note that this mode of dispersal introduces another form of aggregation unrelated to pheromones and not included in our model. Because beetles are picked up and transported together by the wind, the small number of beetles that disperse in this way will be aggregated on the landscape scale.

In addition to the spread speed, we compare the peak fraction of infested trees to an outbreak in Yellowstone studied by Klein et al. (1978). Using our cutoff of 25 cm, or 10 inches to correspond with the units in their analysis, we find the peak fraction of infested trees to be *≈* 33% in 1969. This peak fraction would correspond to an infestation that is just able to spread, around *φ/c* = 0.25, which makes sense given the relatively low density of lodgepole pine in the area. We are also able to estimate *c* from the peak emergence of *≈* 25000 beetles/acre with 27.4 infested stems/acre, or about 900 total beetles per infested tree. This corresponds to *c ≈* 600 assuming a 2/3 sex ratio. Given the peak outbreak fraction, this would correspond to *φ ≈* 150 beetles/tree. All of these parameter estimates are within a reasonable range. More broadly, the relatively high peak fraction for infestation that we find is consistent with observations of very high cumulative pine mortality above 85% in some outbreaks (Hiratsuka et al. 1981; Hiratsuka et al. 1982).

Given the similarity in models, we additionally compare our results to those of Duncan and Powell (2017). Their model also uses the forest age structure from Duncan et al. (2015) together with integrodifference equations to model spread. Their model does not reflect as many aspects of mountain pine beetle biology, in particular it does not include an Allee effect and includes overcompensation, and it uses a Gaussian kernel for dispersal, which does not appear to be as consistent with data as the Laplace kernel (Goodsman et al. 2016; Carroll et al. 2017). However, their model does allow for analytical solutions for spread speed and infestation size. They find lower invasion speeds of only a few hundred meters per year, even for very high population growth rates, and lower peak infestations of around 10% of total trees. The lower spread speed also points to the importance of including long distance dispersal. We note that the smaller fraction of infestation may result from differences in our definition of peak infestation. Overall, their approach allows for much more detailed analytical work and finds results consistent with their data from Sawtooth National Recreation Area, whereas our approach includes additional biological mechanism at the cost of analytical solutions and is consistent with data as outlined above.

The results from the spread into higher resistance trees and into the buffer zone indicate that more resistant trees could potentially slow or stop local spread and thus may give managers more chance to respond. However, these results should be treated with caution given that our model does not include long distance dispersal (see above) or the effects of variable dispersal rates with forest density (Powell and Bentz 2014; Powell et al. 2018). These effects may cause beetles to simply disperse beyond a resistant buffer of moderate size, and therefore small buffers have the potential to accelerate beetles spread on larger spatial scales. With that caveat, our results indicate that the value of host resistance required to stop spread is well within the plausible range for forests, around *φ* = 350 or *φ/c* = 0.27, and small changes in host resistance can substantially reduce local spread speed given it is roughly linear below this threshold (Fig. 4). From a management perspective, these results suggest that lodgepole pine breeding programs selecting for host resistance may be effective in slowing local spread given the heritability of host resistance (Yanchuk et al. 2008; Six et al. 2018). Of course, this would require substantial investment in breeding programs as well as making it a priority to reforest with MPB resistant trees. Even if such a program were prioritized, it would likely take decades before its effects were observed as these resistant trees would have to mature. Additionally, if resistant trees were planted only on a small spatial scale, beetles may be able to disperse beyond resistant stands. We also note that because beetle brood size depends on factors other than host resistance, most importantly weather (Safranyik and Wilson 2006), a particularly good year for beetles may reduce the effect of planted resistant trees. This may be especially true with climate change, which may increase host susceptibility on a similar time scale to the implementation of such programs (Régnière and Bentz 2007; Safranyik and Wilson 2006; Matthews et al. 2018). Note that it is not yet clear how genetic host resistance affects MPB susceptibility mechanistically (Six et al. 2018; Cullingham et al. 2020), and so here genetically resistant trees could have an increased threshold *φ*, a decreased brood productivity *c*, or some combination of both.

In addition to pointing towards the potential success of breeding programs, our results have implications for MPB spread into the boreal forest and into jack pine. There is some evidence that lodgepole pine hosts in regions where MPB has not been found historically are less resistant (Cudmore et al. 2010). If this is the case, we might expect that spread through the remaining lodgepole pine in Alberta will progress more quickly and be more severe. MPB range expansion into the jack pine and the boreal forest is uncertain. Jack pine may have lower thresholds for defense as it is less able to produce toxic resin, but MPB may also have reduced reproductive potential in jack pine (Safranyik and Linton 1982; Cerezke 1995; Safranyik et al. 2010; Rosenberger et al. 2017a; Rosenberger et al. 2017b; Bleiker et al. 2023). More research is needed to determine a biologically plausible range of the ratio *φ/c* in this novel host.

In addition to simplifying assumptions around dispersal, our model does not include several important biological features. Here we have assumed that all susceptible trees produce the same number of beetles within a susceptible stand, and are equally susceptible. We interpret this as an average over the successfully attacked trees in a stand assuming a reasonable distribution of stand diameters. A more realistic model might more explicitly include mean stand dbh to reflect that larger trees both have a higher threshold for defense and once infested can produce many more beetles (Safranyik and Wilson 2006). These parameters also vary with climate variables, as for example cold winters will kill larvae and reduce brood size, and drought will reduce tree thresholds (Régnière and Bentz 2007; Safranyik and Wilson 2006; Matthews et al. 2018). Connecting these parameters more directly to climate would also allow for a more detailed analysis of the effect of climate change on pine beetle spread. Finally, we do not model the different behaviour of the beetle with density (Safranyik and Carroll 2006). Beetles in the endemic state preferentially attack weaker host trees until their population grows to the point where they are able to mass attack and overcome the defenses of large, more vigorous trees (Safranyik and Carroll 2006; Boone et al. 2011; Bleiker et al. 2014). In order to model this behaviour, we would have to include low vigor trees in our model.

Overall, our model predicts transient outbreaks in susceptible stands where beetles are able to infest large fractions of susceptible trees and spread at the speed of a few kilometers per year. We find that the ability for beetles to spread into new stands, and the size of the resulting outbreaks, changes quickly for realistic values of host resistance. More research is needed to determine the susceptibility of novel hosts, especially jack pine. Our simulation results point towards the potential of breeding programs that select for MPB resistant trees to be able to slow or stop local spread. However, for such a program to be successful it would require significant investment in selecting for and prioritizing planting these trees, and it would likely be decades before we would see the effects of such a program.

## Acknowledgements

We would like to thank all Lewis Research Group members, in particular Evan Johnson, Xiaoqi Xie, and Kévan Rastello, for their feedback on this project. We would also like to thank the members of the TRIA-FoR project for their support. MAL gratefully acknowledges the Gilbert and Betty Kennedy Chair in Mathematical Biology. Funding for this research has been provided through grants to the TRIA-FoR Project to MAL from Genome Canada (Project No. 18202) and the Government of Alberta through Genome Alberta (Grant No. L20TF), with contributions from the University of Alberta and fRI Research (Project No. U22004). This work was supported by Mitacs through the Mitacs Accelerate Program, in partnership with fRI Research. MB acknowledges the support of the Natural Sciences and Engineering Research Council of Canada (NSERC), [PDF – 568176 - 2022].

