## Supporting Information A-F for "Mountain pine beetle spread in forests with varying host resistance"

### A Varying the dispersal parameter $\alpha$

In the text we carefully justify our choice of dispersal parameter $\alpha = 0.001$ m$^{-1}$ to be aligned with the scale of local pine beetle spread. However, this value can vary by orders of magnitude depending on which data we use to parameterize dispersal. Fortunately, the exact choice of $\alpha$ can be thought of as scaling the length units, as long as the scale of dispersal is larger than the scale of aggregation. In other words, the choice of the dispersal parameter corresponds to a choice of units for the length scale.

More concretely, the speed scales with $1/\alpha$, and so what appears as 2 km/year in Fig. 4 with $\alpha = 0.001$ m$^{-1}$ corresponds to 200 m/year with $\alpha = 0.01$ m$^{-1}$. To show this, we plot the speed multiplied by the dispersal parameter $\alpha$ in Fig. A1 across the three orders of magnitude of reasonable values we found for $\alpha$.

Note that with $\alpha = 0.1$ m$^{-1}$, we do see a slower speed, a slightly smaller cutoff, and a small horizontal section when the speed corresponds to 16 m/year. This comes from the discreteness of the grid. When the scale of dispersal is comparable to the scale of aggregation, we do start to see the effects of $\alpha$ in a way that cannot be scaled away. This is true for the size of the infestation as well, which we plot in Fig. A2.

Given that the realistic range for dispersal tends to be much larger than the relevant scale of aggregation, these effects are unlikely to effect MPB spread. As a result, changes in $\alpha$ result only in scaling the speed by $1/\alpha$.

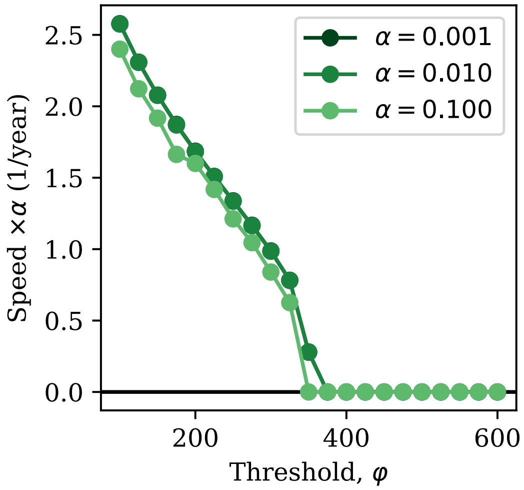


Figure A1: The speed of the transient wave pulse for different values of the host resistance and dispersal distance. As Fig. 4 in the main text, but with different values of the dispersal parameter $\alpha$. The units of $\alpha$ are $m^{-1}$.

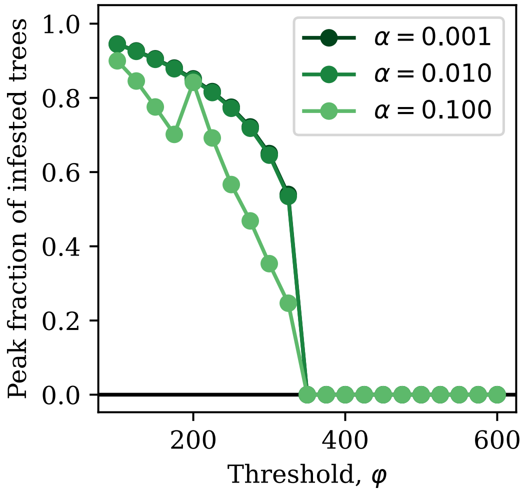


Figure A2: The averaged peak fraction of infested trees in the transient wave pulse for different values of the host resistance and dispersal distance. As Fig. 5 in the main text, but with different values of the dispersal parameter $\alpha$. The units of $\alpha$ are $m^{-1}$.

### B Fitting the aggregation parameter directly to data

While the aggregation parameter $k$ is challenging to connect directly to data, there are a few datasets where we are able to fit beetle attack density distributions with a negative binomial. We here fit attack density distributions to three datasets: Safranyik (1968), Peterman (1974), and Waring and Pitman (1985). With the Peterman (1974) data, we are also able to fit the distribution for the threshold $\varphi$ as the survival fraction of the trees was recorded. All fits were performed by maximizing the log-likelihood.

In Safranyik (1968), 43 infested trees at a Horsethief Creek site and 22 infested trees at an Elk Creek site were felled in 1965 and 1966, both in British Columbia. The attack density on these trees was measured by removing an 8" by 12" bark area at two foot intervals from both the Northern and Southern aspects of the tree up to the highest point of infestation. The attack density reported is per 96 in$^2$ sampling unit. Thus, this dataset combines the attack density within a tree and the attack density across trees. Given the scarcity of available data, we still fit to this site. We show fits for the negative binomial across the two sites in Fig. B1, and combined across the Horsethief and Elk Creek sites in Fig. B2. We additionally fit zero-inflated negative binomials given that they report that approximately 1/4 of the sampling units had zero attack counts. These fits are shown separately in Fig. B3 and together in Fig. B4. Across all fits, $k$ is generally between about 2 and 7, with the zero-inflated negative binomial at Elk Creek giving $k = 26.7$.

In Peterman (1974), attacked trees were sampled in 1973 across two endemic and two epidemic locations, all in British Columbia. Elk Creek and Terrace Creek were epidemic sites with considerable damage and Parson, near Elk Creek, and Lean-to Creek, near Terrace Creek, were endemic sites with small beetles populations and were selected to be similar in geography to the nearby epidemic sites. Attack density was measured by averaging the number of entrance holes across three 1 ft by 1 ft sample areas. In addition to attack density, this study tracked attack success. Attacks were determined to be successful by peeling away bark in up to three random locations on each tree to look for live larvae or pupae. We show fits across all four sites in Fig. B5, and for all data combined in Fig. B6. Here we find $k$ between 6 and 10, except at Elk Creek where $k$ is very large. This could indicate that when beetle abundance is very high, the beetles are able to spread out more uniformly with less aggregation, as beetle pressure was likely highest at this site. However, when we combine the data by endemic and epidemic, the best fit aggregation parameters are very similar (Fig. B7), but it is possible that the Lean-to Creek site is beginning to outbreak when the data was taken (Peterman 1974).

Given that in this dataset, we also have the percent of surviving trees, we can minimize the least squares difference of the survival function and the percent survival $P_S$, $(F(m, k, \varphi) - P_S)^2$ to obtain an estimate for $\phi$ (see Section 2.5). We do this by fixing $m$ and $k$ according to their best fit values from the negative

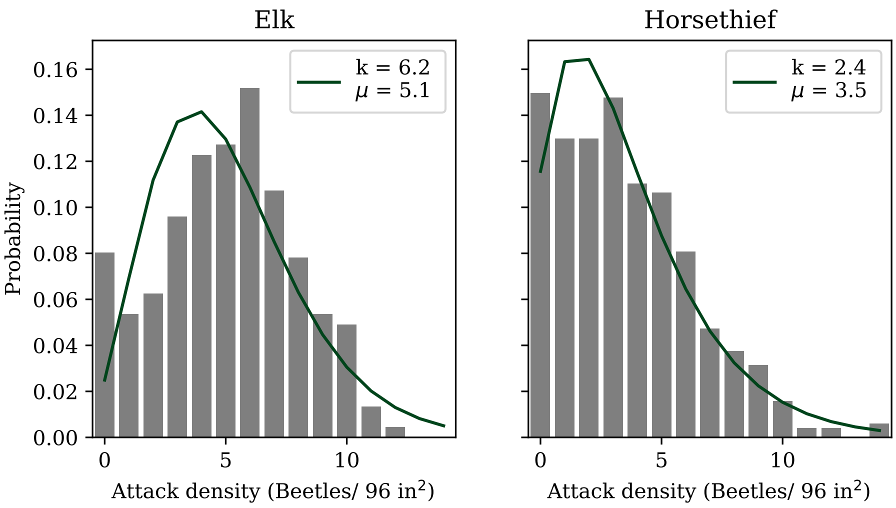


Figure B1: Negative binomial fits to data from 542 sampling units at Horsethief Creek and 604 sampling units at Elk Creek within and across trees.

binomial fit, and then finding the value of $\varphi$ where the survival function is most similar to the true surviving fraction.

In Waring and Pitman (1985), 16 quarter-hectare plots with central circular 200 m$^2$ areas were set up with four different experimental treatments that varied tree vigor. These plots were established in Oregon during an outbreak with high beetle pressure. Each tree with dbh $>$ 10 cm was tagged within the central circle, and the attack density for each tree was measured in 1979. We show the data fit with a negative binomial in Fig. B8, and with a zero-inflated negative binomial in Fig. B9. Note that in this case, there were many attacks with non-zero but very small attack density. For the zero-inflated negative binomial, we set a threshold of 2 attacks per m$^2$, below which we set the attack density to zero. This was mean to give a better fit to the increase of attack densities between 100 and 150 attacks per m$^2$. Fits prefer a smaller $k$ than in the other datasets, which may be the result of more thorough sampling of trees. However, in our model the beetles distribute themselves only among susceptible trees, and so a full census of all trees above 10 cm dbh may be less relevant as we set the cutoff for susceptibility at 25 cm dbh. Ideally then, we would like to know how the beetles distribute themselves among trees with dbh greater than 25 cm, which we cannot fit here.

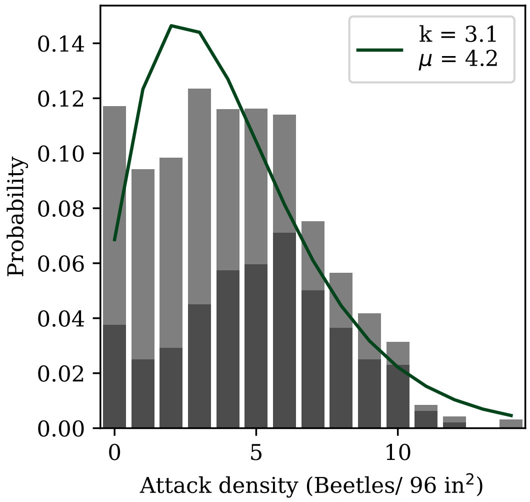


Figure B2: Negative binomial fit to data from all sampling units from both Horsethief Creek and Elk Creek from Safranyik (1968).

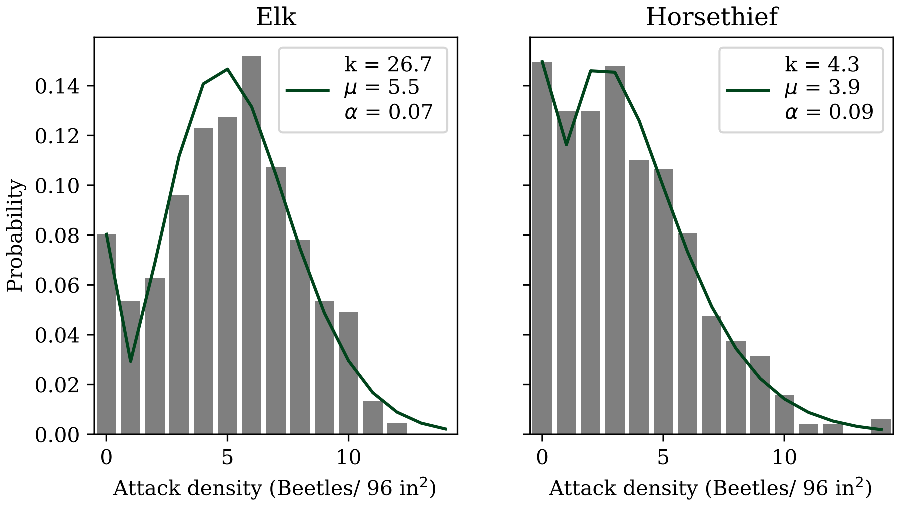


Figure B3: Zero-inflated negative binomial fits to data from 542 sampling units at Horsethief Creek and 604 sampling units at Elk Creek within and across trees from Safranyik (1968).

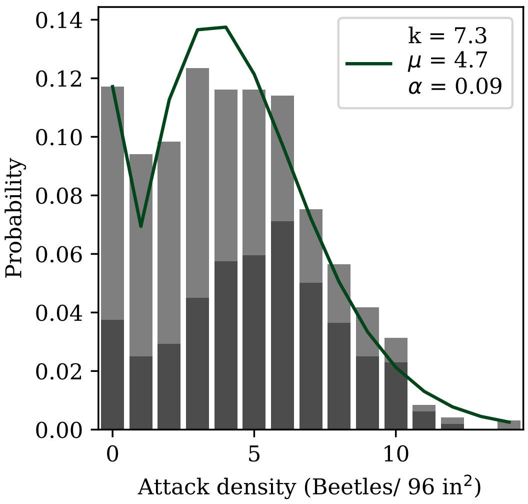


Figure B4: Zero-inflated negative binomial fit to data from all sampling units from both Horsethief Creek and Elk Creek from Safranyik (1968).

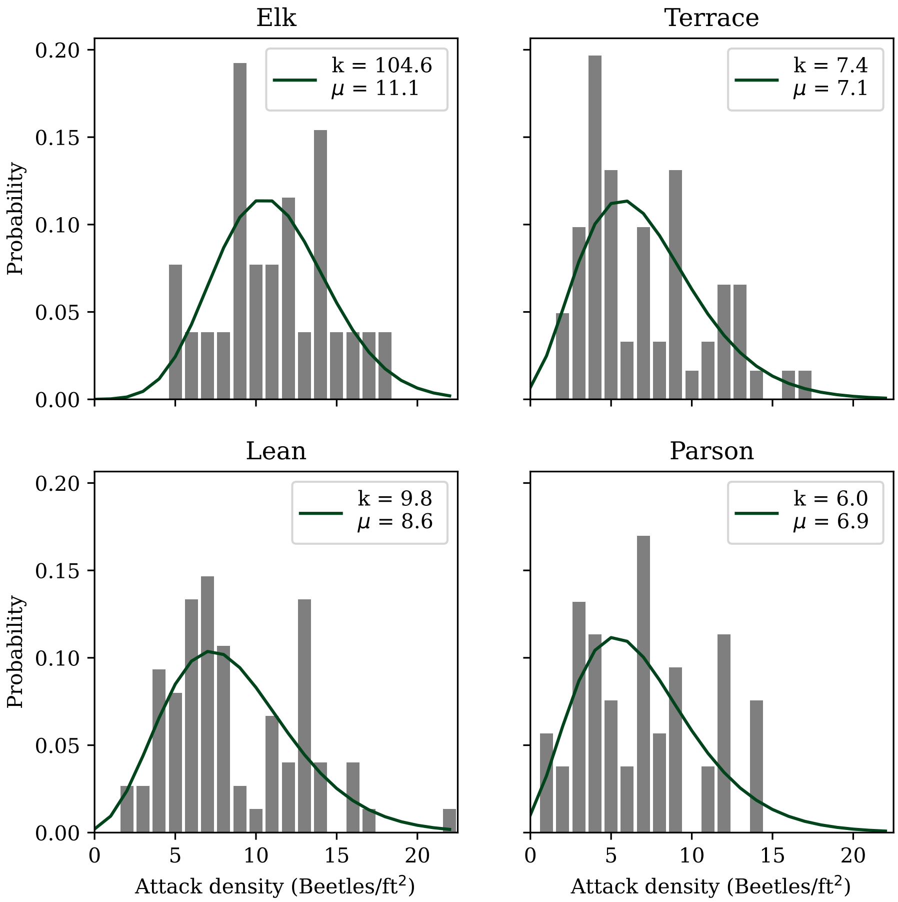


Figure B5: Negative binomial fits to data across four sites from Peterman (1974).

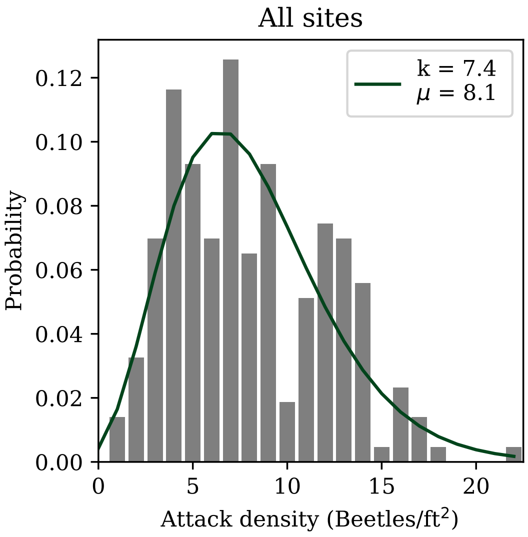


Figure B6: Negative binomial fit to data combined across all four sites from Peterman (1974).

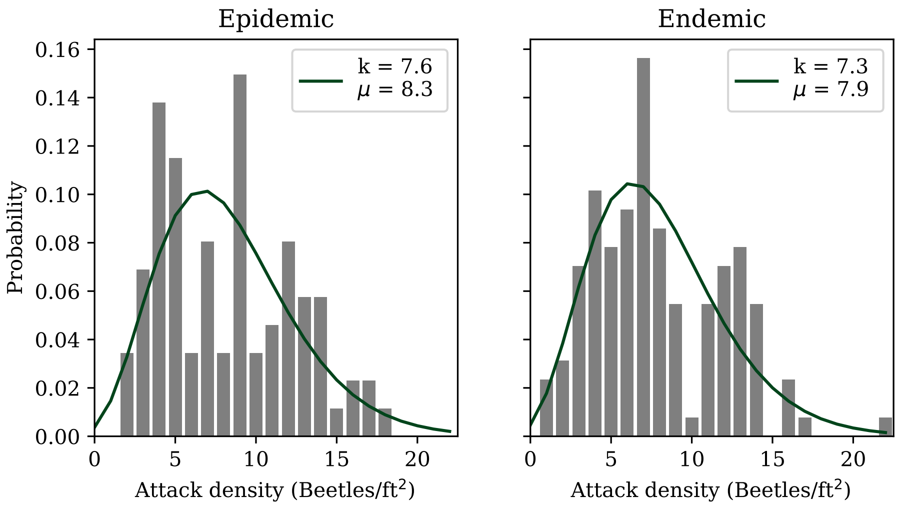


Figure B7: Negative binomial fit to data combined across endemic (Parson and Lean-to Creek) and epidemic (Elk Creek and Terrace Creek) status from Peterman (1974).

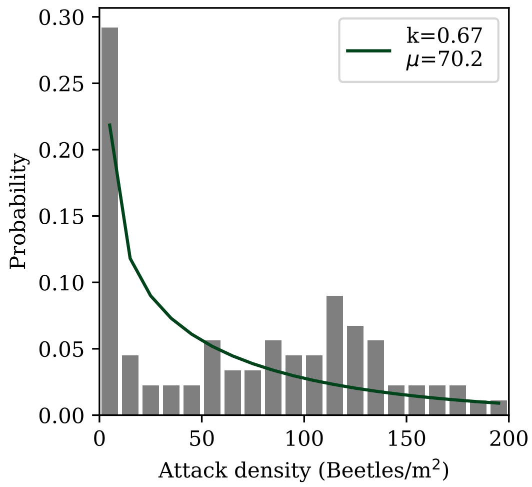


Figure B8: Negative binomial fit to data from Waring and Pitman (1985).

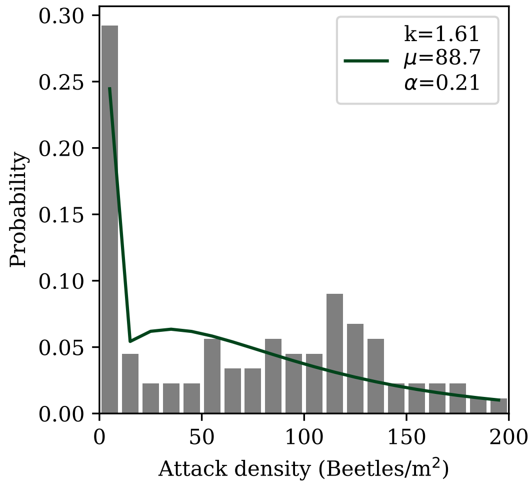


Figure B9: Zero-inflated negative binomial fit to data from Waring and Pitman (1985). All trees with attack density less that 2 attacks per m$^2$ were set to zero for this fit.

### C Varying the cutoffs for the aggregation parameter $k$, $k_{min}$ and $k_{max}$

When we assume that beetles aggregate optimally at the local scale, ie. within a grid cell, we must set a minimum and maximum value for the aggregation parameter $k$ to keep the best fit value finite. This is because at very low numbers of beetles, the beetles would optimally all aggregate on a single tree ($k \to 0$), and at the other extreme when the mean number of beetles per tree is larger than the trees threshold for defense, then the beetles should spread out uniformly ($k \to \infty$). The choice of these cutoffs is very hard to connect to data, as they represent the aggregation at either extreme of density. The cutoff $k_{min}$ corresponds the aggregation of beetles when the mean number of beetles is very small, less than a few beetles per tree, and the cutoff $k_{max}$ corresponds to the aggregation when the mean number of beetles is larger than the threshold, in this case a few hundred or more beetles per tree (Fig. 1).

In the text, we fix these values at $k_{min} = 0.01$ and $k_{max} = 100$. However, note that our justification for $k_{min}$ actually leads to a cutoff that varies with $\varphi$, as the beetle density where the population moves from incipient to epidemic behaviour changes with the threshold $\varphi$. Figure C1 shows the values of $k_{min}$ we find for varying $\varphi$, which vary roughly between 0.007 and 0.008 with a mean of 0.0077. Figure C2 shows how the Allee threshold itself changes when we fix $k_{min} = 0.01$ versus allow it to vary with $\varphi$.

More broadly, given the uncertainty in the cutoffs of both $k_{min}$ and $k_{max}$, we now consider how changes in these parameters would affect our results. We plot the optimized function $F$ in Fig. C3 for 4 orders of magnitude of $k_{min}$ and $k_{max}$, where the first panel shows the function at small values of $m$ to make the difference more apparent. We note that changes in $k_{min}$ can substantially change the Allee threshold, which we plot in Fig. C4. Change in $k_{max}$ do not substantially change the Allee threshold, but instead change the shape of the curve when the mean number of beetles is slightly greater than tree defense threshold. If $k_{max}$ is small enough, it can reduce the upper equilibrium number of beetles (not seen here).

The choice of these cutoffs can affect the dynamics and speed of spread. We plot the speed of the wave for different values of the cutoffs in Fig. C5 and the size in Fig. C6. Increasing $k_{min}$ slightly decreases the speed of spread, and decreases the threshold when the wave is able to advance. In other words, when beetles aggregate less well at low densities they are less able to spread. As long as the wave is able to advance, the peak size of the infestation is similar. Decreasing $k_{min}$ below 0.01 and changing $k_{max}$ have small effects on the speed. We find that the peak size of the infestation decreases slightly with decreasing $k_{max}$.

Note that the values of $k_{min}$ and $k_{max}$ in reality are likely to change with the scale of the grid cell. For example, if the grid cells are larger than the scale of the pheromone production, than the beetles will not be able to aggregate optimally and we should choose a larger cutoff for $k_{min}$.

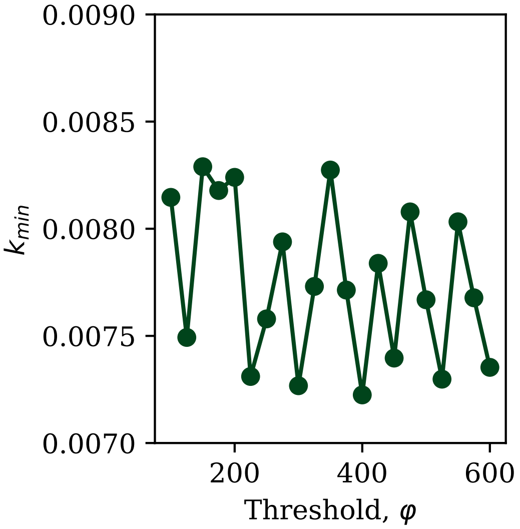


Figure C1: The value of the cutoff $k_{min}$ with varying threshold $\varphi$. This is set by fixing $k_{min}$ when the beetle population moves from incipient to epidemic behaviour, which we here fix at $m = 20\varphi/1225$ (see main text).

Overall, although order of magnitude changes to these values can alter the exact values for the speed or size of spread, the qualitative conclusions from the main text will remain similar. As in the main text, we note that a full treatment of $k_{min}$ would require a model for beetle behaviour at low population densities, which we leave for a future study.

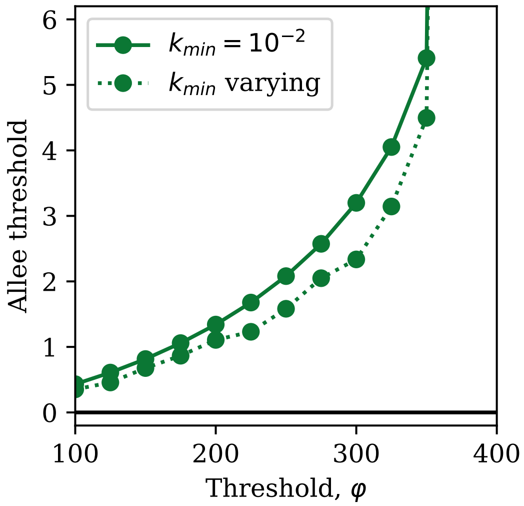


Figure C2: The Allee threshold for varying $k_{min}$ with $\varphi$ compared to fixing $k_{min} = 0.01$.

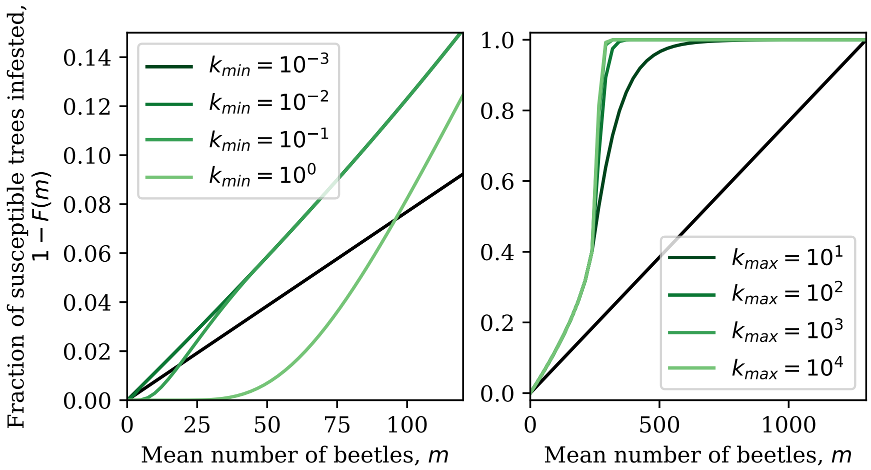


Figure C3: The optimized function $1-F(m)$ representing the fraction of susceptible trees infested for different values of the cutoffs for $k_{min}$ and $k_{max}$. Note the different x-axis ranges between the two plots, as changing $k_{min}$ has a negligible effect for larger values of $m$. We fix the threshold at $\varphi = 250$ and $c = 1300$.

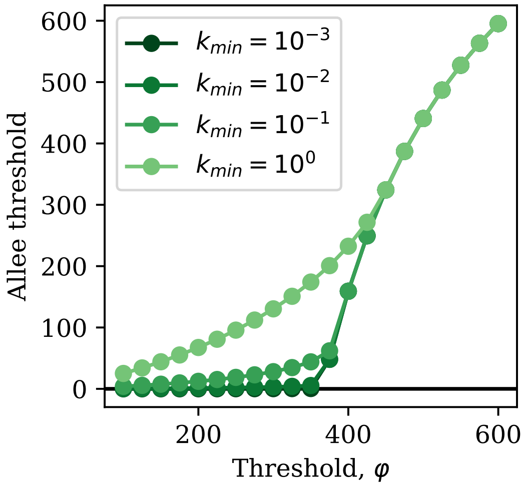


Figure C4: The Allee threshold for varying orders of magnitude in the cutoff $k_{min}$.

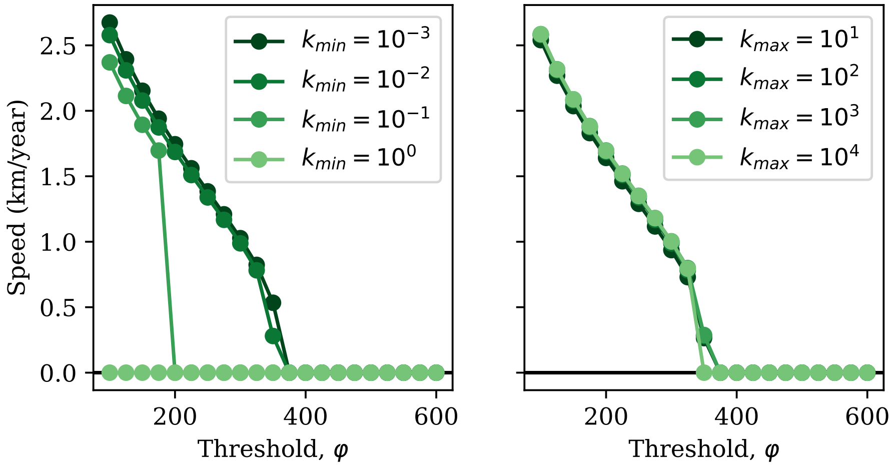


Figure C5: The speed of the transient wave pulse for different values of the host resistance and aggregation cutoffs. As Fig. 4 in the main text, but with different cutoffs $k_{min}$ and $k_{max}$.

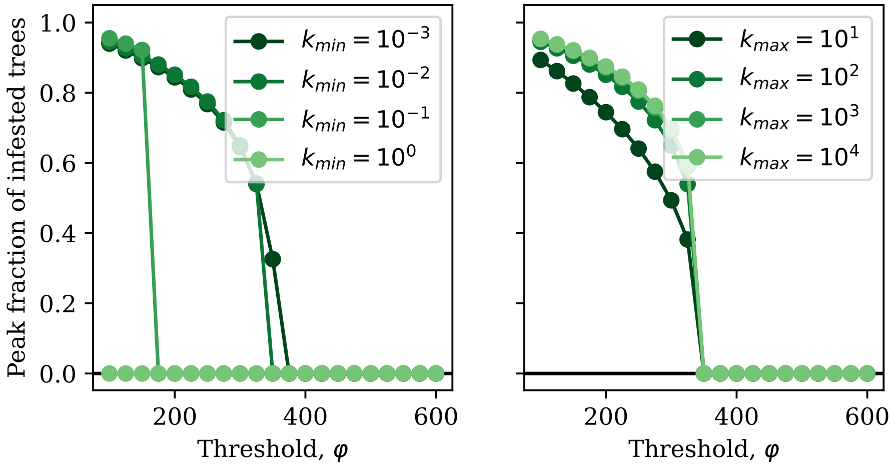


Figure C6: The averaged peak fraction of infested trees in the transient wave pulse for different values of the host resistance and aggregation cutoffs. As Fig. 5 in the main text, but with different cutoffs $k_{min}$ and $k_{max}$.

### D Additional information on parameter estimates

In the main text, we summarize our estimates for the parameters for population dynamics. We here provide more detail about the data and methods used to obtain these estimates. The population dynamics parameters are the number of years before trees become susceptible $N$, the juvenile survival $s$, the brood size parameter $c$, and the threshold parameter $\varphi$. For $N$ and $s$, the parameters relating to forest growth and survival, we will assume susceptible trees are lodgepole pine.

#### D.1 Age classes of juvenile trees $N$

As in the main text, we note that we define susceptible trees as trees that produce more beetles than it takes to overcome their defenses. These trees are sources (rather than sinks) for beetles. This definition is necessary as we assume that brood size is the same across all susceptible trees, when in reality it varies strongly with tree diameter (Safranyik and Wilson 2006). This means that small trees that are attacked generally produce very few beetles, and a stand composed entirely of small trees would not support an outbreak population.

We estimate the number of age classes of juvenile trees $N$ using two methods. We estimate it directly when tree age was recorded in outbreak data, and we estimate it by finding a dbh cutoff for susceptibility and combining that with growth estimates. To estimate this parameter directly from data, Fig. 8 in Safranyik and Wilson (2006) (adapted from Safranyik (1968)) shows that the youngest attacked tree in a stand in Southeastern BC is around 40 years old, with many more trees attacked around age 55. Another direct estimate comes comes from a report by Shore and Safranyik (1992). This report assigns susceptibility to MPB attack to stands depending on overall stand age, and defines low susceptibility as stands under 60 years old, intermediate susceptibility as stands between 60 and 80 years old, and high susceptibility as stands over 80 years old.

We now turn to the second method. We begin by estimating the dbh cutoff for susceptibility. In an outbreak in Yellowstone, nearly all trees below 7 inches in diameter ($\approx$ 18 cm) survived (Klein et al. 1978), and so our cutoff should be above 18 cm. An outbreak in Oregon only appeared to grow when trees above 23 cm remained in the stand (Mitchell and Preisler 1991), which could indicate trees above that dbh are beetle sources. There is additional evidence supporting a similar cutoff and suggesting that trees up to 25 cm in diameter are beetles sinks and above that are beetle sources (Safranyik et al. 1974; Safranyik and Wilson 2006). Another study found that this cutoff is likely to be higher, between 12 and 14 inches ($\approx$30 to 35 cm) (Cole and Amman 1969). Given this range, we set the dbh cutoff at 25 cm.

We now estimate the radial growth rate. The average growth rate found in a mixed stand of lodgepole pine and Douglas fir after MPB thinning is less than 2 mm per year, with a maximum around 2 mm per year immediately after thinning (Heath and Alfaro 1990). At another site in BC, the average radial growth rate for lodgepole pine between 3 and 23 years old was 2.4 mm per year, where the stands are subject to some controlled thinning. For older lodgepole pine in Alberta, Chhin et al. (2008) find a mean radial growth rate of 1.24 mm. We note that the MPB model of Duncan et al. (2015), where our model for forest growth comes from, uses this method as well. They estimate the dbh cutoff as 20 cm and assume 2 mm per year for radial growth as the maximum growth rate from Heath and Alfaro (1990), which gives $N = 50$. Given that we want to know the radial growth rate of juvenile trees, these data point towards a reasonable range between 1.5 and 2 mm per year, which gives $N = 63$ to 83 with a dbh cutoff of 25 cm.

Rather than using radial growth, we can use forestry site index estimates together with a height to dbh model to obtain an approximate age when stands will reach 25 cm in diameter. The site index is defined to be the height the average tree will reach 50 years after reaching breast height. We can therefore use the site index to estimate the height of trees in different regions, and then convert the height to dbh using the model of Huang (1999). We find that a site index of 18 to 21 m gives around 50 years for the tree to grow from breast height to 25 cm dbh. Given that it takes about 6 years in BC or Alberta for a tree to reach breast height (Huang et al. 1994; Thrower et al. 1994; Nigh and Love 1999), this means that for a site index of 18 to 21 an approximate estimate would be $N = 56$. From Monserud et al. (2008), the site index of the lower foothills of Alberta is roughly 18 to 21 m. It is lower elsewhere in the range and generally higher in BC (Thrower et al. 1994). Overall then, $N$ depends on the area under consideration, but can reasonably vary between 50 and 80. We set $N = 60$ in simulations.

#### D.2 Survival fraction of juvenile trees $s$

In British Columbia, where lodgepole pine is particularly well adapted, experiments studying different methods of forest site preparation find between 90 and 97% of juveniles survive after 10 years, without a large effect from different site preparation techniques (Bedford and Sutton 2000). This corresponds to $s = 0.99$ to 0.997. In Alberta, there is evidence for lower survival rates. Across different microsites, lodgepole pine survival was found to be between 60% and 85% after 7 or 8 years (Brown and Navratil 1995), corresponding to $s = 0.94$ to 0.98. In another study, 5, 10, and 15 year survival rates were measured for lodgepole, hybrid lodgepole x jack, and jack pine (Rweyongeza et al. 2007). Survival was found to be similar across all species. We estimate $s$ across years from this data and find $s = 0.98$ for 5 year survival, $s = 0.984$ for 10 year survival, and $s = 0.987$ for 15 year survival. This study finds that mortality is higher when the tree is younger, which is why we find $s$ increases as we consider a longer time interval. Given this, we would expect $s$ to be even larger as it assumes juvenile survival is the same across all years until the juveniles become susceptible. From this data, we set $s = 0.99$ with a reasonable range from $s = 0.98$ to 0.997.

#### D.3 Tree surface area

For the remaining two parameters, $c$ and $\varphi$, we have to estimate the surface area of a tree attacked by beetles to convert between units of per m$^2$ and per tree. From the data in Klein et al. (1978), we know most beetle attacks occur on the bottom 6.9 m of a tree. We can then take the weighted average of the diameters of the trees in the stand, excluding the 6 inch diameter class, and we find that the average tree is 9.75 inches ($\approx$25 cm) in diameter. This corresponds to about 5.5 m$^2$ per tree of attacked surface area. Other estimates for the surface area come from Powell and Bentz (2009), who assume 6.25 m of the tree is attacked and assume an average of 32 cm dbh to obtain 6.3 m$^2$ per tree, and Strohm et al. (2013) who assume 7.5 m of the tree is attacked and use the diameters from data to obtain 4.4 to 8.1 m$^2$ per tree. We will assume 6 m$^2$ per tree is attacked if we lack stand size data, but use direct stand size data where possible.

#### D.4 Brood size $c$

The brood size per tree $c$ is likely to be highly variable depending on the year and tree given that it is affected by weather, in particular overwinter temperatures and drought conditions, and tree size and vigor (Safranyik and Wilson 2006). Additionally, this parameter will increase with the number of beetles successfully overcoming tree defenses, which we do not account for in our model. This makes estimates of this parameter particularly uncertain, and so we attempt to estimate it from as many sources as possible. We first note that as in the main text, we assume 2/3 of all beetle offspring are female (Amman and Cole 1983; Safranyik and Wilson 2006). This is important as our model tracks only the female beetle population dynamics.

In several studies, this parameter can be estimated by multiplying the number of offspring per attack, the number of attacks per m$^2$, the surface area attacked, and the sex ratio of the offspring. Using this method, Goodsman et al. (2016) (which provides the basis for the beetle population dynamics in this model) estimate $c$ to be 537 female beetles per tree. They assume the optimal beetle attack density of 62 beetles per m$^2$ found by Raffa and Berryman (1983) and scale up to 606 beetles per tree using the Klein et al. (1978) attack data as their stand size distribution is similar. They then multiply by 2/3 to obtain the number of female beetles and finally multiplying by the number of beetle offspring per entry hole at their site, 1.33.

We here use this method to obtain estimates from other data sources. In the attack data from an outbreak in Yellowstone (Klein et al. 1978), the average number of offspring per entry hole was found to be 2.2, though this is highly variable, and attack densities where found to range between 50 and 100 beetles per m$^2$. Using the data directly from this study that 5.5 m$^2$ per tree is attacked (see above), this corresponds to between 400 and 800 female beetles per tree. Another study found that between 8 and 11 pupae survived per attack with attack densities between 40 and 80 beetles per m$^2$, with the peak at 60 beetles/$m^2$ (Raffa and Berryman 1983). Assuming 6 m$^2$ per tree is attacked, and that all pupae survive to adulthood, this corresponds to about $c = 1300$ to 3000 female beetles per tree. Observations in the Sawtooth National Recreation Area find between 1300 and 1500 emerging beetles per $m^2$ and approximately 6.3 $m^2$ attacked per tree (Powell and Bentz 2009), which corresponds to between 5000 and 6300 female beetles per tree.

This parameter can also be estimated directly from data of brood size per tree (Cole and Amman 1969; Safranyik 1968; Safranyik and Wilson 2006; Powell and Bentz 2009). Fig. 8 in Safranyik and Wilson (2006) shows that trees produce between a few hundred and 12 000 beetles, depending heavily on tree size, and provides a linear fit to dbh. We take that fit and average over the stand size distribution Klein et al. (1978) to obtain a stand average of about 1250 female beetles per tree. A similar range of brood production was found by Cole and Amman (1969), who find between 300 and 15 000 beetles emerge per tree depending on tree size. Fitting a phenology and population growth model, Powell and Bentz (2009) also find between 10 000 and 20 000 beetles per tree depending on the temperature of the phloem. Given all of this, we assume a reasonable range of $c$ to be from 400 to 6000 female beetles per tree, with the upper range only attained during large outbreaks. We fix the parameter at $c = 1300$ female beetles per tree for simulations.

#### D.5 Host resistance $\varphi$

The most common estimate in the literature for the threshold $\varphi$ is to take the value of 40 beetles per $m^2$ from aggregation manipulation experiments in Raffa and Berryman (1983) and scale it as needed (Powell and Bentz 2009; Strohm et al. 2013). Assuming 6 $m^2$ per tree as above, this corresponds to $\varphi = 240$ beetles per tree. Data for the range of the threshold against host vigor comes from Waring and Pitman (1985), with a maximum likelihood fit by Lewis et al. (2010) finding a range from 25-120 beetles per $m^2$ needed for attack success. This corresponds to a range of $\varphi = 150$ to 720 beetles per tree, though note the upper range is likely only attained in a few trees rather than at the stand level. The mean threshold for hosts in this data is 54 beetles per $m^2$, or 324 beetles per tree.

We can also estimate this parameter by fitting the attack density data from Peterman (1974) (see Supporting Information B and find the attack threshold varies from 41 beetles per $m^2$ to 97 beetles per $m^2$ across four sites, or 60 beetles per $m^2$ when the data is combined across sites. Using 6 $m^2$ attacked per tree, this corresponds to $\varphi = 250$ to 520 beetles per tree across sites or 360 beetles per tree combining the data. We assume a reasonable range of thresholds to be from 150 to 600 and fix $\varphi = 250$ when needed.

### E Wave speed in a simplified model

Numerically, we find that the wave is no longer able to advance for values of the host resistance per tree brood $\phi/c \approx 0.27$. Given the complicated age structure model of the forest and the Allee effect, we are not able to calculate this threshold value analytically (though we can estimate it as in the main text). We can obtain a rigorous upper bound for this value if we remove forest structure from the model and additionally assume that trees are not killed after infestation. This is equivalent to setting $N = 0$ and all trees as susceptible, and assuming infested trees are immediately replaced by new susceptible trees in the following year. We will refer to this model in this section as the simplified model. In this case, Li and Otto (2022) prove that for an integrodifference equation with an Allee effect, the sign of the speed is zero if and only if $\int_0^c dm(1 - F(m) - m/c) = 0$. Technically, this is only true if $1 - F(c) = 1$ as we have to scale by the upper equilibrium, but we can see this is approximately true from Fig. 1.

We plot the value of the integral $\int_0^c dm(1 - F(m) - m/c) = 0$ for varying threshold $\phi$ with fixed $c = 1300$ in Fig. E1. Note that we cannot compute this integral analytically as we optimize the aggregation $k$ for different values of the threshold $\varphi$. We find numerically the integral crosses 0 when $\varphi \approx 750$, or for $\varphi/c \approx 0.58$.

In addition to computing the value of the integral, we run simulations as in the main text with this simplified model to see how the speed changes with $\varphi$ away from this threshold value. We plot the speed of these simulations in Fig. E2 and find it is generally an upper bound for the speed in the complete structured forest model. Note that the value of $\varphi$ where the simplified model with can no longer advance (ie. when $\int_0^c dm(1-F(m)-m/c) = 0$) is similar to the value of $\varphi$ when the wave in the complete model reverses direction (Fig. E2). This makes sense as it occurs roughly when $\int_0^c dm(1 - F(m) - m/c) < 0$, at which point the upper stable equilibrium no longer exists (Brush and Lewis 2023) and the beetle population disappears (in both models).

We note that the speed of spread in the complete model appears to always be slower than in the simplified model. Biologically, this makes sense as susceptible trees are always replaced immediately in the simplified model, given more fuel for the beetles. This means it takes less time for the outbreak to reach peak density and thus it will spread more quickly.

Another mathematically tractable estimate for the speed in the case of no forest structure and immediate tree replacement comes from assuming a caricature Allee effect, where

$$1 - F(m) = \begin{cases} 0, & m < m_{allee} \\ 1, & m \geq m_{allee}, \end{cases} \tag{E1}$$

where $m_{allee}$ is the value of the Allee threshold. In this case, the speed can be calculated as $-\log(2 * m_{allee}/c)/\alpha$ (Lutscher 2019). This is zero when the Allee threshold is half of the brood size parameter, or around 650. This corresponds to roughly $\varphi = 650$ (extrapolating from Fig. 2), though below this the speed is

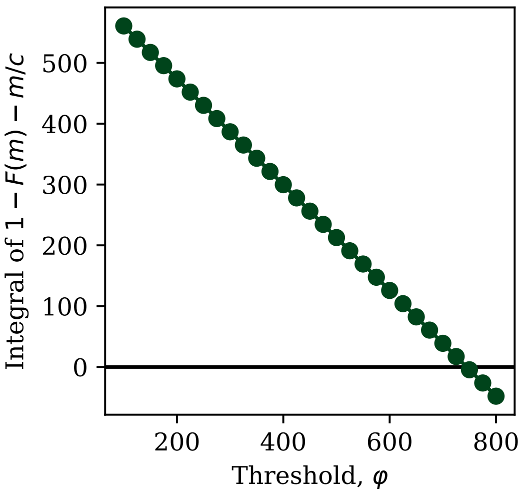


Figure E1: The integral of $1 - F(m) - m/c$ with $c = 1300$. The sign of this integral should determine the sign of the wave in the simplified model.

dramatically overestimated (Fig. E3).

Overall, then, we are able to approximately determine the value of the threshold where the wave reverses direction as $\varphi/c \approx 0.58$, which is well above the reasonable range during an outbreak. Further analytical work may be able to determine the approximate threshold value for when the speed of the wave is 0.

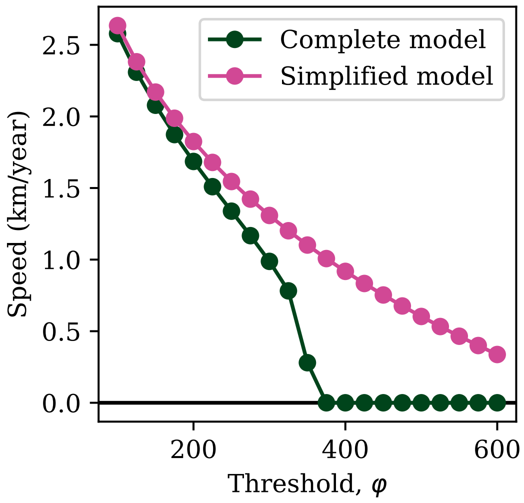


Figure E2: The speed of the wave from Fig. 4 compared to the speed of the wave in the simplified model, where there is no forest structure and infested trees are immediately replaced, but there is still an Allee effect.

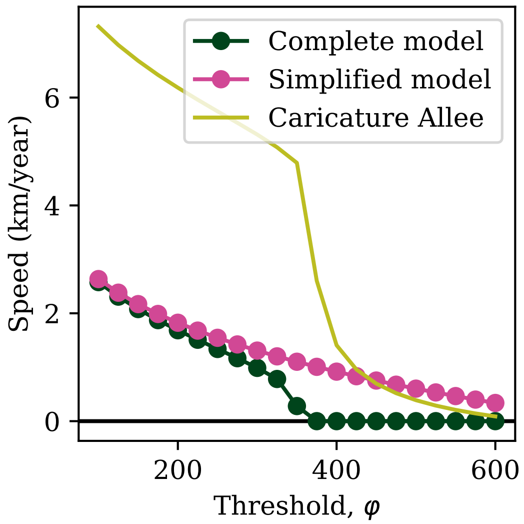


Figure E3: The speed of the wave from Fig. 4 compared to the speed of the wave in the simplified model, as well as the speed of the wave with a caricature Allee effect.

### F Varying the grid size

In the simulations in the main text, we set the grid size at $\Delta x = 16$ m according to the estimated relevant scale of beetle aggregation. However, we note that this grid size may be too small in stands with lower density as it will correspond to only a few stems within a cell. This would invalidate our model assumption about a large number of trees in a stand (Brush and Lewis 2023). In particular, stand densities below about 400 stems/ha correspond to fewer than 10 stems within a cell. In these cases, it may be more appropriate to select a grid size of 32 m, or even 64 m. These distances would still be mostly consistent with data about the range of pheromones, but would lead to more suitable number of host trees within a cell. With that said, we show here that as long as the scale of dispersal is substantially larger than the size of the grid, the exact values for the grid size should not significantly affect our results.

We additionally investigate how large the scale of the grid must be compared to the scale of dispersal in order to see wave pinning. This happens when the wave is no longer able to advance because of the discreteness of the grid in systems with an Allee effect (Keitt et al. 2001). Because we want to compare the grid size and the scale of dispersal, the relevant dimensionless parameter is $\alpha \cdot \Delta x$. In the main text, this is equal to $0.001 \cdot 16 = 0.016$. Given that this is still very small, we will instead consider $\alpha = 0.1$ m$^{-1}$ and $\alpha = 0.01$ m$^{-1}$, roughly corresponding to results from Safranyik et al. (1992) and Robertson et al. (2007), respectively. We then simulate wave spread as outline in the methods in the main text and calculate the speed for various values of the grid size $\Delta x$.

Figure F1 shows the results of these simulations. Note that the speed changes only slightly for most values of the grid size, until the wave becomes pinned. . With the very short dispersal parameter $\alpha = 0.1$ m$^{-1}$ observed in the mark recapture experiment of Safranyik et al. (1992), we start to see pinning when the grid spacing is $2^5 = 32$ m, which is a realistic upper bound for the scale of aggregation. With $\alpha = 0.01$ m$^{-1}$, we only see pinning when the grid spacing is $2^8 = 256$ m, which is well beyond a reasonable scale of pheromone mediated aggregation. We therefore conclude that pinning due to the Allee effect is unlikely to be relevant for MPB. Further, we find that for realistic values of grid size according to pheromone dispersal distances (16, 32, and 64 m) the speed of beetle spread is roughly equivalent, even when the scale of beetle dispersal is an order of magnitude shorter than considered in the main text.

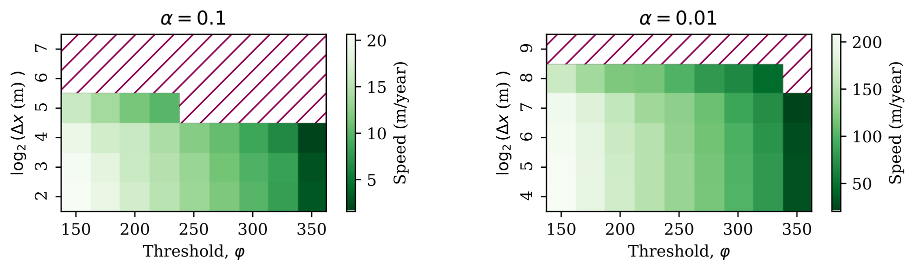


Figure F1: The effect of the scale of aggregation (the grid size) on the speed of spread. When beetles aggregate over very large scales, comparable to their scale of dispersal, the wave can become pinned (indicated by regions with dark red hashes). However, this scale for aggregation is biologically unrealistic.
